# Cerebral organoids recapitulate interneuron chain migration in the human cortex

**DOI:** 10.64898/2026.09.30.729488

**Authors:** Sakurako Nagumo Wong, Mark Alan Noble, Michael Fischer, Leo Meyer, Maria Novatchkova, Ema Novakova, Denise van der Heijden, Lucas Barea Moya, Susana Gonzalez-Granero, Jose Manuel Garcia-Verdugo, Sara Merino-Aceituno, Nina S. Corsini, Juergen A. Knoblich

**Affiliations:** Institute of Molecular Biotechnology of the Austrian Academy of Sciences (IMBA); Vienna, Austria; Vienna BioCenter (VBC), Doctoral School of the University of Vienna and Medical University of Vienna; Vienna, Austria; Department of Mathematics and Scientific Computing, University of Graz; Graz, Austria; Faculty of Mathematics, University of Vienna; Vienna, Austria; Laboratory of Comparative Neurobiology, Cavanilles Institute of Biodiversity and Evolutionary Biology, University of Valencia and CIBERNED-ISCIII; Valencia, Spain; Research Group in Immunotherapy and Biomodels for Autoimmunity, Health Research Institute La Fe; Department of Neurology, Hospital Universitario y Politécnico La Fe; Valencia, Spain; Biomedical & Tissue Engineering Laboratory (BTELAB), Fundación de Investigación del Hospital General Universitario de Valencia (FIHGUV); Valencia, Spain; Department of Neurology, Medical University of Vienna; Vienna, Austria

## Abstract

The human brain is incomplete at birth, with streams of late-born interneurons migrating into the postnatal cortex. This process is absent in rodents, limiting mechanistic investigation to post-mortem tissue. Here, we show that human cerebral organoids cultured for over six months recapitulate postnatal interneuron migration, enabling its mechanistic and genetic dissection. Live imaging and mathematical modeling revealed that interneurons intrinsically migrate in chains, while astrocyte interactions promote stream organization. Spatial and single-cell transcriptomics identified a distinct molecular signature of human postnatal migratory streams enriched for neurodevelopmental disorder-associated genes. Perturbation of the autism-associated genes *CNTN5* and *FGF14* differentially disrupted chain organization and migration. Thus, long-term cerebral organoids enable mechanistic analysis of previously inaccessible aspects of postnatal human brain development and their contributions to neurodevelopmental disease.

## Main Text

The coordinated arrival of GABAergic interneurons (INs) from the subpallial ganglionic eminences (GEs) and integration throughout the developing cortex are critical for establishing cortical connectivity during late gestation(1). Unlike in rodents, the protracted development of the human brain has resulted in multiple prominent migratory streams that emerge late in gestation and persist after birth, continuing to supply immature neurons to cortical regions for months to years postnatally. The medial migratory stream (MMS) and Arc-derived neurons are present postnatally in gyrencephalic mammals, while the entorhinal cortex (EC) stream persists up to 11 months after birth in humans but not macaques(2–5). The evolutionary expansion of the CGE and an increased proportion of CGE-derived inhibitory neurons in humans(6, 7) suggests that their protracted migration to the cortex may be important for higher order cognitive functions. Postnatal streams of CGE-derived INs are most prominent several months after birth, within the critical window of network plasticity when infants receive external sensory input from their surroundings(8, 9). The protraction of these processes may also pose additional risks for neurodevelopmental disorders (NDDs) in humans, including autism spectrum disorder (ASD), schizophrenia, and epilepsy(10–13). However, as peri- and postnatal tissue is difficult to obtain and can only provide static snapshots, there is a crucial need for tractable model systems in which the dynamics, genetics, and function of these processes can be interrogated.

Cerebral organoids(14–17) and other brain organoid models(18–20) have emerged as a developmental model capable of recapitulating the major stages of human brain development. They can be maintained as long-term cultures(21, 22) and continue to follow transcriptomic and epigenetic trajectories of human brain development *in vivo* even after 250 days in culture(23, 24). Here, we reveal that long-term cultured cerebral organoids recapitulate the formation of late born IN streams of CGE origin, allowing mechanistic and genetic investigation of their unique mode of chain migration. We quantify their migration dynamics in real time and identify distinct transcriptional networks and disease associations that define this unique late-stage neurodevelopmental process.

### Late-born interneurons in CGE-enriched organoids form compact streams

In humans, migratory streams become progressively composed of CGE INs during the first few months after birth(3, 5), aligning with the later migration and integration of CGE INs in the developing cortex(6, 7). To model this, we modified a ventral forebrain patterning protocol to enrich for CGE inhibitory neurons, using a previously described(25) *Dlxi5/6-eGFP* reporter introduced into three hiPSC lines (Fig. 1A, fig. S1A). Reducing the duration of ventral patterning compared to our previous protocol(25) yielded organoids with pronounced expression of the CGE marker COUP-TFII (Fig. 1B). At day 25, these organoids (referred to as CGE-enriched organoids, CGE-Os) exhibited delayed *Dlxi5/6-eGFP* expression compared to MGE-enriched organoids (fig. S1B), aligning with the later *in vivo* development of CGE INs compared to MGE(26, 27). By two months, CGE-Os showed abundant Dlx5/6+ neurons marked by calretinin (CALB2) and secretagogin (SCGN), a CGE- and dorsal LGE-derived cortical IN marker arising from mid-gestation in humans(28, 29) (Fig. 1C). Organoids showed abundant expression of the forebrain marker FOXG1, IN marker DLX2, and COUP-TFII, with minimal expression of NKX2-1 and the excitatory marker TBR1, demonstrating robust telencephalic CGE lineage specification (fig. S1C).

**Fig. 1.**
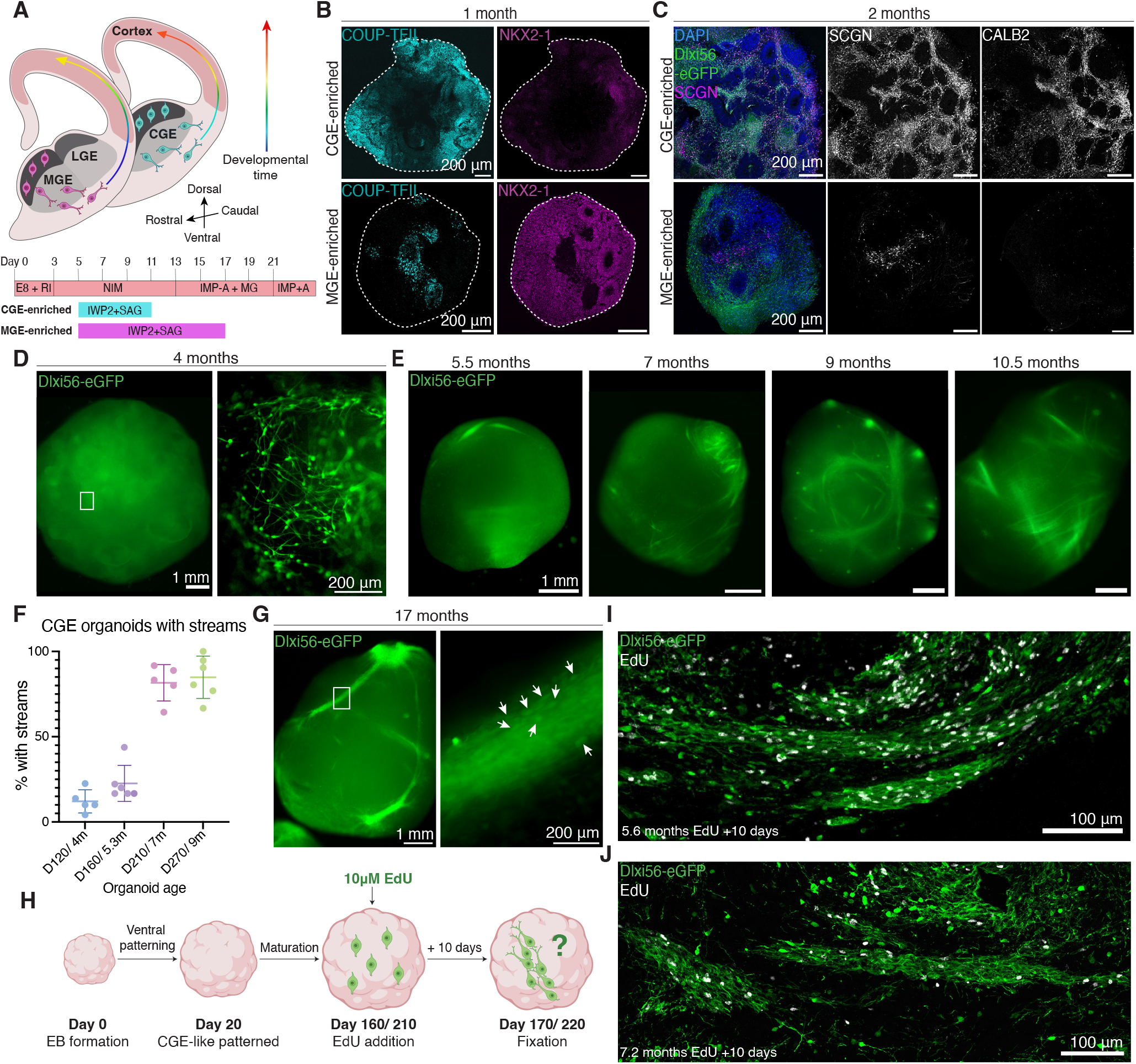
Late-born INs in CGE-enriched organoids form compact streams. **(A)** Top: A schematic of medial, lateral, and caudal ganglionic eminences (MGE, LGE, CGE), with MGE INs (magenta) migrating first to the cortex, followed by CGE INs (cyan) migrating later in development (arrows). Bottom: CGE-enriched and MGE-enriched organoid patterning timelines. **(B)** Immunostainings at 1 month show high COUP-TFII (cyan) and low NKX2-1 (magenta) in CGE-enriched organoids and vice versa in MGE-enriched.**(C)** Immunostainings at 2 months show co-expression of Dlxi56-eGFP with SCGN and CALB2 in CGE-enriched organoids (top) and minimal co-expression in MGE-enriched (bottom). **(D)** Widefield GFP images of 4-month CGE-enriched organoid (left) and zoom-in (right) shows dispersed IN distribution. **(E)** Widefield GFP images of whole CGE-enriched organoids across 5.5-10.5 months in culture, showing the timeline of Dlxi56-eGFP+ IN stream progression. **(F)** Quantification of percentage of organoids with streams per batch across organoid age in 3 cell lines. Each point represents 1 organoid batch. Mean +/-SD = 4m: 9.32 +/-4.85, 5.3m: 22.66 +/-10.58, 7m: 81.66+/-10.67, 9m: 84.88+/-12.42 SD. **(G)** Left: Widefield GFP image of whole CGE-like organoid at 17 months. Right: Individual Dlxi56-eGFP+ INs are oriented along the stream (white arrows). **(H)** Schematic of EdU birth-dating with 2h pulse EdU. **(I)** and **(J)** Immunostainings of Dlxi56-eGFP streams show EdU+ cells in white at **(I)** 5.6 months and **(J)** 7.2 months.

At 4 months, CGE-Os had diffuse GFP expression with individual INs exhibiting elongated migratory morphology (Fig. 1D). Strikingly, from 5.5 months on, CGE-Os formed condensed stream-like structures of GFP+ INs that became more prevalent with organoid age (Fig. 1E). By 7 months, an average of 81.7% of CGE-Os across batches exhibited streams (Fig. 1F, fig. S1D), while MGE-enriched organoids had minimal stream formation (fig. S1E). GFP+ streams were still prevalent in CGE-Os at 17 months, a postnatal equivalent timepoint (Fig. 1G). Organoid stream formation therefore reflects the late gestation to postnatal timeline in humans(3, 5). One defining feature is their emergence from continuously proliferating late-born neuronal progenitors(5). To determine whether the GFP+ streams are composed of late-born neurons, birth-dating experiments were performed using a 2-hour pulse of 5-ethynyl-2’-deoxyuridine (EdU) in 5- and 7-month CGE-Os (Fig. 1H), timepoints after the proliferation peak at 1-2 months(30). Ten days after EdU addition at 5 months, EdU+ nuclei were abundant within GFP+ INs streams (Fig. 1I). At 7 months, newly generated EdU+ cells still incorporated into GFP+ streams (Fig. 1J, fig. S1F). These findings demonstrate that CGE-Os continuously give rise to late-born GFP+ INs that contribute to condensed stream-like structures at late stages, reflecting the developmental changes of the human CGE subventricular zone (SVZ).

### CGE interneuron streams in organoids exhibit chain migration

*In vivo*, postnatal neuronal migration is directed from the SVZ to distinct cortical regions, where neurons then disperse to integrate into cortical circuits(3, 5). To assess for directed migration *in vitro*, CGE-Os were fused with age-matched dorsal organoids at 4 months, slightly before streams arise (fig. S2A–C). 7 days after fusion, INs had started to migrate from the CGE-enriched into the dorsal organoid (fig. S2D). At a perinatal-equivalent timepoint of 9 months, late-fused CGE-Dorsal assembloids exhibited defined streams that were dispersed throughout the assembloid (fig. S2E).

To deduce whether postnatal-equivalent GFP+ streams were composed of CGE INs, immunostaining was performed on 13-month assembloids (Fig. 2A). Analyses of GFP+ streams in both assembloids and single CGE-Os confirmed their primarily CGE IN identity, with abundant co-localization of GFP with SCGN and CALB2, as well as high COUP-TFII+ and minimal NKX2-1+ expression across cell lines (Fig. 2A-C, fig. S3A,B). GFP+ INs expressed both doublecortin (DCX) and Polysialic acid-neural cell adhesion molecule (PSA-NCAM), commonly expressed in migratory neuroblasts, alongside few KI67+ dividing cells (Fig. 2B, fig. S3A). Streams did not express TBR1 or the oligodendrocyte progenitor cell (OPC) marker OLIG2 (fig. S3A). Streams identified earlier at 6 months were also highly COUP-TFII+ (fig. S3D), already suggesting a bias towards CGE identity at this time point.

**Fig. 2.**
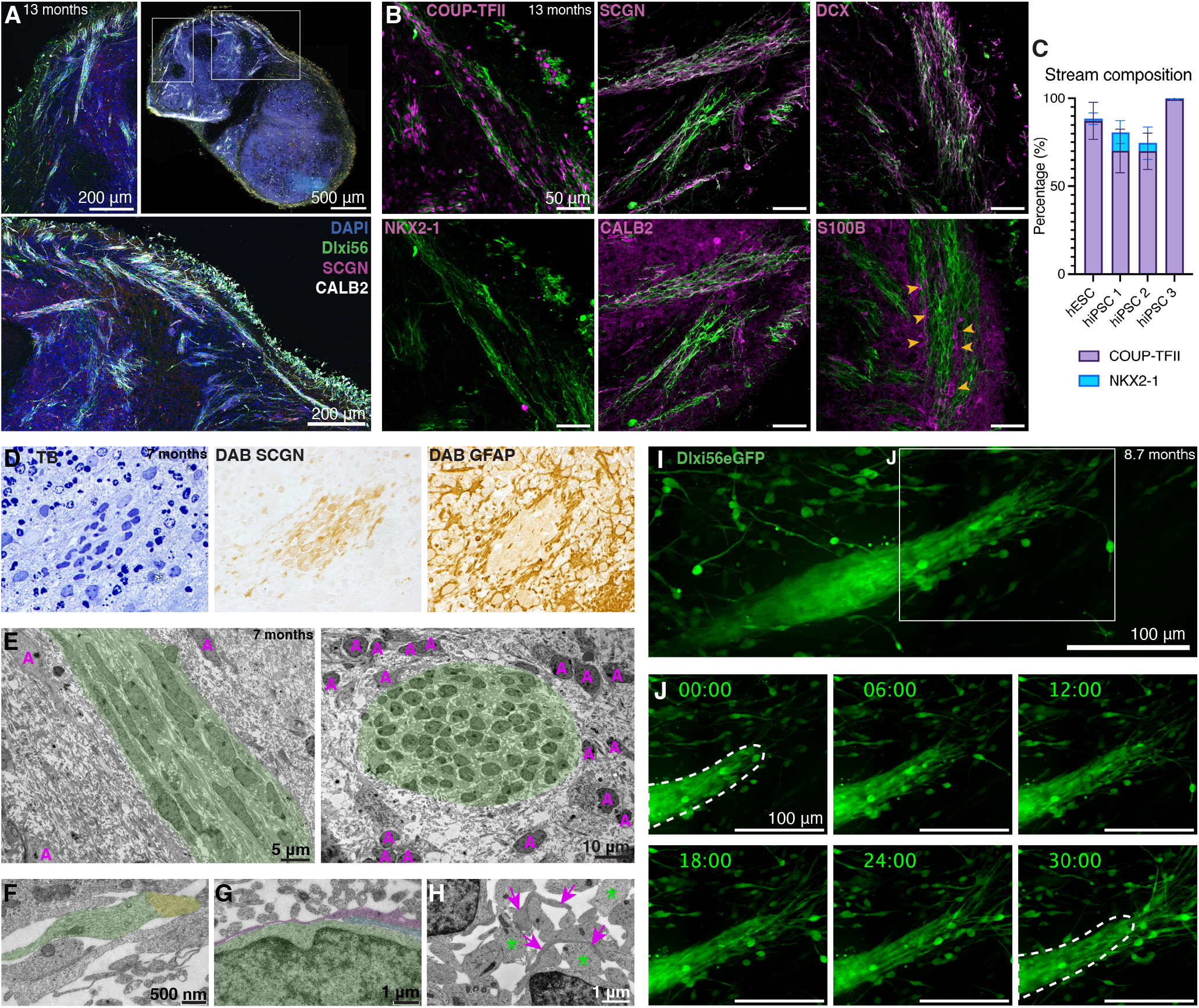
CGE IN streams in organoids exhibit chain migration. **(A)** Immunostainings of a CGE-D assembloid at 13 months. Whole section (top right) with zoomed-in regions (left, bottom) show Dlxi56-eGFP IN streams marked by SCGN and CALB2. From hiPSC1. **(B)** Immunostaining in 13-month organoids (all markers in magenta). Streams are positive for COUP-TFII, SCGN, CALB2 and DCX and are negative for NKX2-1 and are surrounded by S100B+ glia (yellow arrows). **(C)** Percentage of COUP-TFII+ CGE versus NKX2-1+ MGE cells normalized to all DAPI+ cells in streams. Data from 3-14 streams from 3-4 organoids across 4 cell lines. Error bars mark SD. **(D)** Toluidine blue (TB) marks nuclei and DAB stainings of SCGN (middle) and GFAP (right) on 7-month organoid streams. **(E)** Electron microscopy images of longitudinal-(left) and cross-sections (right) of 7-month organoid streams. EM of 6 organoids from hiPSC line 3. Migratory streams (green) are surrounded by astrocytes (magenta A). **(F)** IN leading process (green) containing microtubules and growth cones (yellow). **(G)** IN soma (green) within stream, with another IN process (blue) surrounded by a thinner astrocytic process (magenta). **(H)** Thin, dark astrocytic processes (magenta arrows) are distinguishable from larger lighter IN processes (green *). **(I)** Live time-lapse imaging of Dlxi56-eGFP+ streams in whole organoids at 8.7 months. **(J)** Time-lapse stills at t=0, 6, 12, 18, 24, 30 hours of imaging, focusing on the leading edge outlined in **(I)**. Dotted white line marks leading edge at t=0.

We observed that streams were surrounded by S100B+ (Fig. 2B) and GFAP+ glial cells, with GFAP+ processes infiltrating the streams (fig. S3A,C). Cross-sectional 3, 3’-diaminobenzidine (DAB) staining confirmed that SCGN+ cells were encapsulated by GFAP+ astrocytes (Fig. 2D), recapitulating glial tubes that surround postnatal migratory streams *in vivo*, providing structural support and molecular guidance (3, 31). Ultrastructural electron microscopy analysis revealed chains of connected immature neurons with elongated nuclei and multiple nucleoli(32) demonstrating features of migratory morphology, and surrounded by astrocytes with larger, lighter nuclei(3, 33–35) (Fig. 2E). These features are strikingly similar to what has been described for the Arc and EC stream(3, 5). Immature neurons within streams had abundant mitochondria (fig. S3E) and leading processes contained microtubules and growth cones (Fig. 2F), markers of active, directed migration. Astrocytes in the surrounding neuropil were identifiable by formation of gap junctions and the presence of glycogen, visualized as electrolucent spaces (fig. S3F). Tracing the expansions from astrocyte soma revealed thin, dark processes compared to the thicker, lighter microtubule-rich processes of migratory neurons (fig. S3G). This distinction allowed us to observe astrocytic processes both surrounding the chain and directly interacting with neuronal processes within streams (Fig. 2G,H). Astrocytes formed tripartite synapses outside of the streams (fig. S3H), indicating physiological function in late-organoid cultures.

We next performed live time-lapse imaging on 9-month whole organoids to visualize the migratory behavior of INs within streams (Fig. 2I, Supplementary Video 1). Strikingly, GFP+ INs exhibited dynamic movement with neurons sliding over one another, forming a connected chain that led to directional advancement at the leading edge (Fig. 2J). This collective migration was highly reminiscent of chain migration in the postnatal rostral migratory stream (RMS) (32, 33, 36). Together, these findings establish organoid streams as predominantly CGE-derived and astrocyte-encapsulated, enabling the live mechanistic investigation of this perinatal process in a human-specific *in vitro* environment.

### Late-born CGE interneurons exhibit an intrinsic propensity for collective migration

During embryonic brain development, INs tangentially migrate to the cortex along defined paths, while exhibiting independent, exploratory behavior on a single neuron level(37). This migration mode has been replicated in ∼2–3-month cerebral organoids(16, 25, 38, 39) and resembles INs in young CGE-Os. This is in contrast to the connected chains of INs that emerge in the third trimester and persist postnatally, which is observed in >6-month CGE-Os. As INs in streams are late-born, they may follow a distinct intrinsic program. Alternatively, the cell-type milieu and extracellular matrix develop over time both in organoids and the human brain(23, 40–42). We therefore asked whether the transition from dispersed to chain migration reflects an intrinsic property of late-born neurons or arises from the aging cellular environment.

For this purpose, we developed a 2D explant culture system by dissecting and plating organoids on Matrigel (Fig. 3A). In accordance with transcriptomic(41, 43) and epigenetic(44, 45) comparisons to human fetal datasets combined with onset of stream formation, we refer to organoids <4 months in culture without streams as ‘early’ and organoids over 5.5 months as ‘late’. After two weeks of plating, early-organoid INs from 2–4-month explants migrated out in a dispersed manner (fig. S4A,B), with COUP-TFII immunostaining confirming their CGE identity (fig. S4C). In contrast, late-organoid INs from 6-month explants maintained chain-like structures in their outward migration after two weeks of plating, forming extensive, interconnected clusters (Fig. 3B). This was not seen in dorsally patterned explants, which instead yielded projections (fig. S4D). Widefield images taken just one day after plating showed that the initial outward migration from late explants immediately occurred in connected chains (fig. S4E,E’), resembling neuroblasts migrating out of postnatal mouse SVZ explants(32). Immunostaining showed that late-migrating INs were SCGN+ (fig. S4F).

**Fig. 3.**
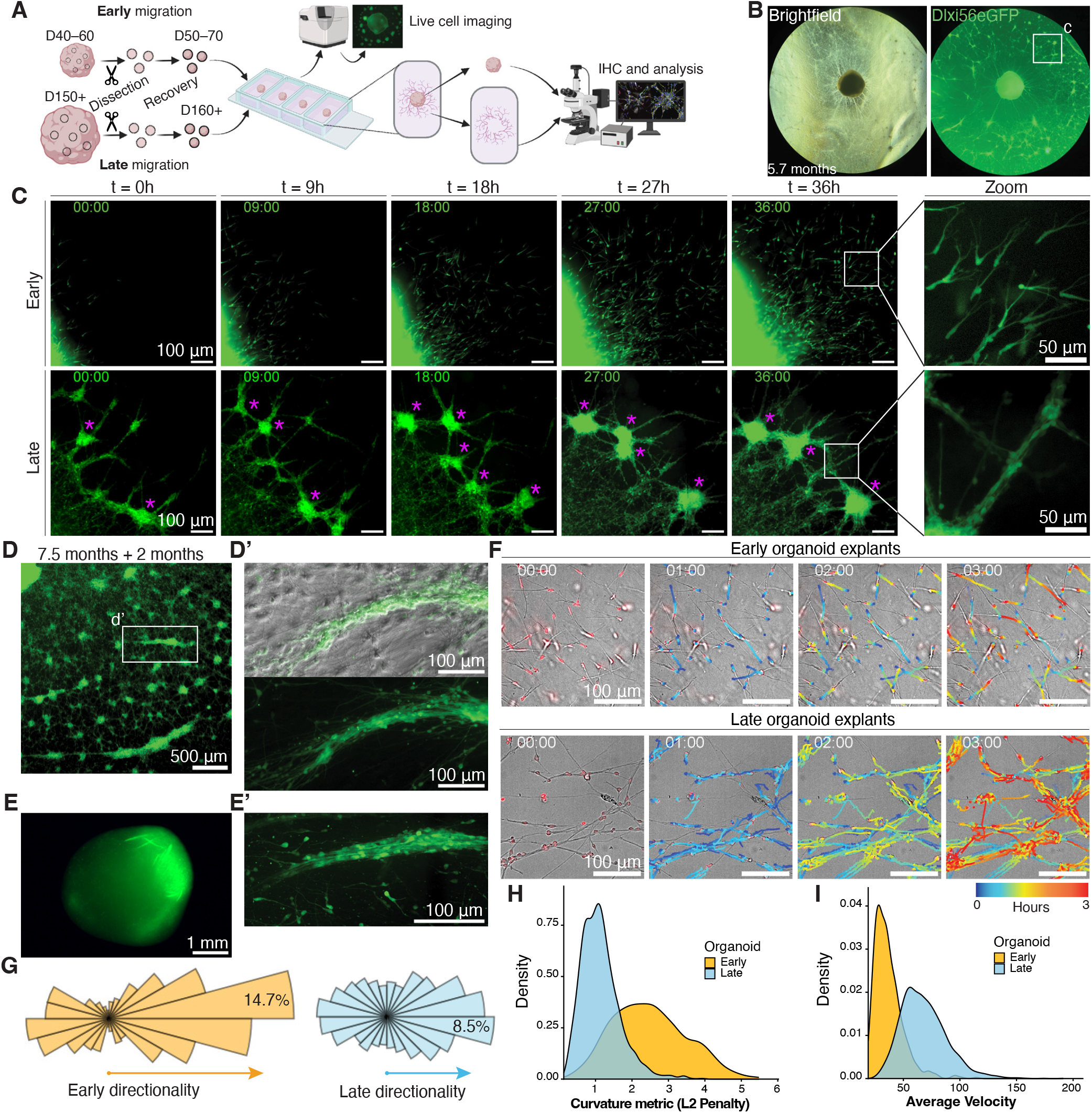
Late-born CGE INs exhibit an intrinsic propensity for collective migration. **(A)** Schematic of 2D explant assay. Organoids are dissected, recovered for >10 days, then plated in chambers. After >5 days after plating, migrated cells are time-lapse imaged and fixed for further analysis. **(B)** Widefield overview images of plated D160 organoids and migrated cells in brightfield (left) and GFP (right) after 2 weeks at D174/5.7 months. White square represents zoom-in in (c), hiPSC line 1. **(C)** Time-lapse imaging stills of Dlxi56-eGFP INs migrating out of organoid explants. Images across 36h from D40/ 1.3-month organoids (top) and D160/ 5.3-month organoids (bottom). Zoom-ins show differences in IN organization in early vs. late (right). Specifically in late explants, INs form dense clustered regions (magenta*). **(D)** Widefield GFP image of a late organoid explant plated at D227/ 7.5 months and imaged 2 months after, with IN clusters fusing linearly. **(D’)**, Zoom-in of fused clusters that resemble streams in brightfield+GFP (top) and GFP only (bottom). **(E)** Streams in whole 6-month CGE organoid. **(E’)** Zoom-in of whole organoid stream, closely resembling (d’). **(F)** Tracks formed over a 3-hour period analyzed with Trackmate software in early (top) and late (bottom) videos. **(G)** Polar plots showing directionality of all IN tracks from early vs. late time-lapses. 14.7% of INs migrate along a main axis away from early explants while only 8.5% follow a main axis in late explants. N=6 experiments. **(H)** Density plot showing distribution of track curvature (L2 penalty), comparing early and late videos. A lower L2 penalty represents a more linear/ straight track. N=6. **(I)** Density plot showing distribution of velocity, comparing early and late videos. N=6.

We next compared early- and late-explant migration dynamics with time-lapse imaging performed at the leading edge (Fig. 3C). This revealed striking differences in their dynamics: early-organoid INs exhibited dispersed, saltatory migration with general directionality away from the explant, whereas late-organoid INs formed a network of chains connecting cell clusters that dynamically fused to form larger clusters (Fig. 3C, Supplementary Videos 2,3). Two months after plating, multiple neuronal clusters had fused linearly to form dense GFP+ IN streams in several regions (Fig. 3D,D’), resembling IN streams in whole organoids (Fig. 3E,E’). These findings suggest that late-organoid INs have an intrinsic propensity to associate during migration, allowing chain formation to be recapitulated outside of the organoid environment.

To quantify the differences in early- and late-organoid migration dynamics, tracking was performed on explant time-lapses imaged in 2-minute intervals (Fig. 3F, fig. S4G,H). This first revealed differences in overall migration directionality, with early migration characterized by strong global directionality away from the explant (Fig. 3G). In contrast, late-organoid INs migrated within an interconnected network of chains oriented in multiple directions, resulting in lower global directionality. We next assessed individual track straightness by penalizing curvature in migratory tracks (Fig. 3H, fig. S4I,J, Methods). Early-organoid INs followed less constrained trajectories, resulting in greater track curvature and higher penalty values, while late-organoid INs exhibited straighter trajectories within chains and therefore lower values.

Chain migration is characteristic of one of the longest migration routes in brain development(46). Consequently, it is significantly faster than other migration modes(47). Indeed, INs from late organoids migrated faster than those from early organoids, with a mean velocity of 68 *µ*m/h compared to 37 *µ*m/h respectively, with velocity going up to 140 *µ*m/h in late conditions (Fig. 3I, fig. S4K, Supplementary Videos 4,5). Notably, chain migration speed in the mouse RMS is 70-80 *µ*m/h(47) and can increase to 130 *µ*m/h(48, 49), similar to our chain-migrating INs. Together, our data show that late-migrating INs exhibit an intrinsic propensity for collective migration, forming chains with more directed trajectories and higher velocities than early-migrating INs.

### Interneuron and astrocyte interactions direct stream organization

One consistent observation in late explants was the formation of dense IN clusters (Fig. 3C*). We considered whether this was a passive consequence of high IN density or driven by interactions with other cell types. Both *in vivo*(33) and in late CGE-Os, astrocytes closely interact with chain-migrating INs, raising the question of whether similar interactions were maintained in explants.

Immunostaining of early 2-month explants confirmed that early-born INs migrated in a dispersed manner and independent of GFAP+ astrocytes (Fig. 4A). In late 8-month explants, INs initially migrated independently of GFAP+ cells but formed connected chains within 5 days of plating, supporting an intrinsic late migratory state (Fig. 4B). However, 2 weeks after plating, GFAP+ processes were found to interact with INs at dense cluster regions (Fig. 4C, fig. S5A). GFAP+ cells displayed characteristic astrocyte morphology, and GFP+ INs were observed along astrocytic processes (Fig. 4D, fig. S5B). Time-lapse imaging one month after plating revealed dense astrocyte-IN clusters alongside astrocyte-sparse regions where INs continued to migrate in chains (fig. S5C, Supplementary Videos 6,7). These regional differences suggested that local astrocyte interactions may progressively influence IN organization as they chain migrate from the explant.

**Fig. 4.**
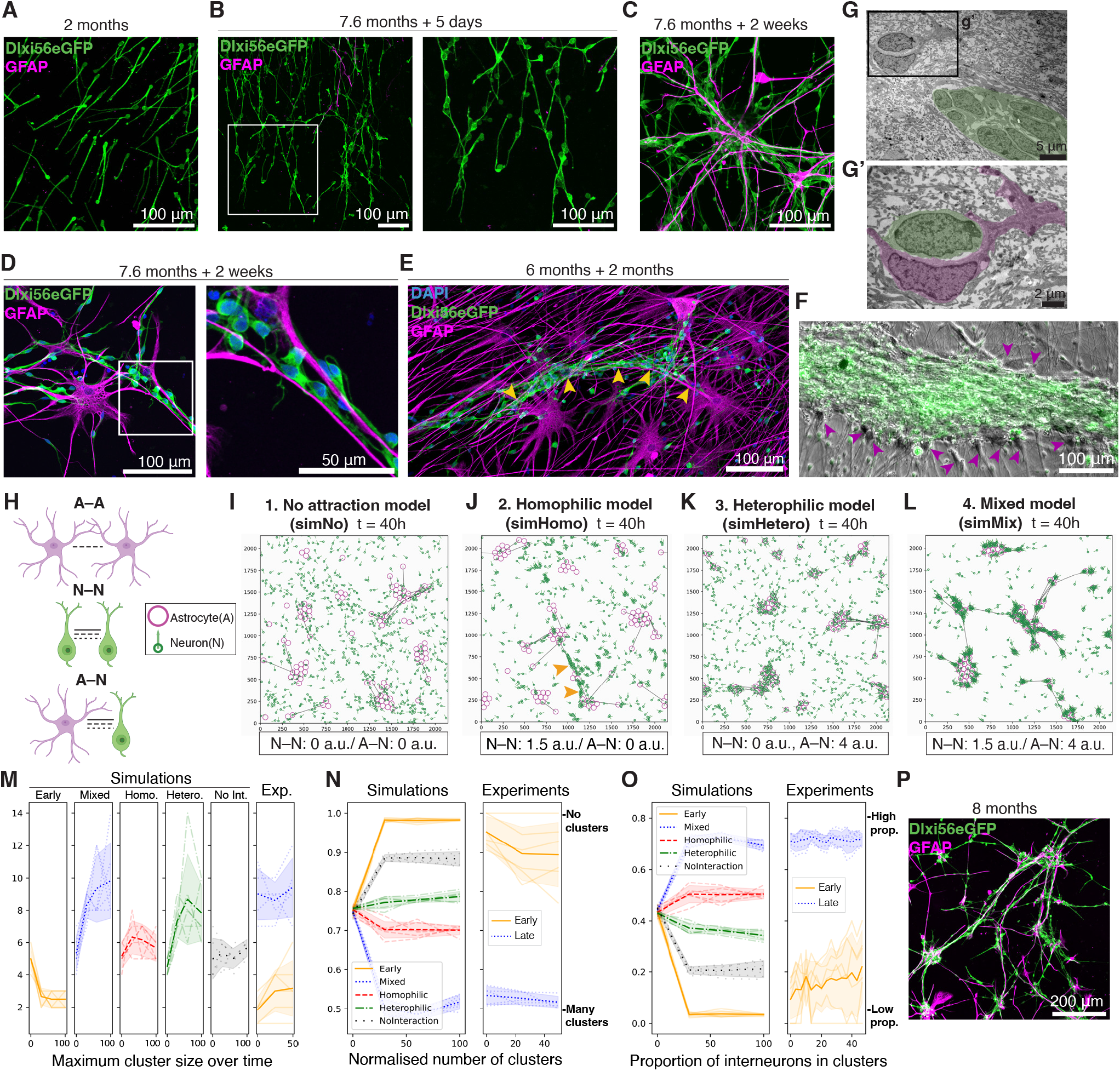
Interneuron and astrocyte interactions lead to stream organization. **(A)** Immunostaining of 2-month explants show INs migrate in a dispersed manner and independently of GFAP+ cells. **(B)** In D230/ 7.6-month explants, INs migrate in chains but independent of GFAP+ cells (magenta) 5-days post plating. **(C)** After 2 weeks of plating, INs in 7.6-month explants interact with GFAP+ processes (magenta). **(D)** Immunostaining at D245/ 7.6 months show GFAP+ cells have clear astrocyte morphology, and INs migrate along astrocytic processes. **(E)** Immunostaining of long-term explant at 246 days/ 8-month organoid, 2-months plated. GFP INs form a chain and are surrounded by GFAP+ astrocytes (magenta), migrating along astrocytic processes (yellow arrows). **(F)** Overlay image of brightfield and GFP from live imaging of thicker stream in long-term explant (8-month organoid, 2-months plated) show unlabeled cells lining GFP+ streams. **(G)** Electron microscopy image of stream (green) in a 7-month whole organoid. **(G’)** Zoom-in from (g) shows an astrocyte (magenta) encapsulating an IN (green) close to the stream. **(H)** Schematic of astrocyte-astrocyte (A-A), neuron-neuron (N-N) and A-N (astrocyte-neuron) interactions used in the mathematical model, where neuron=interneuron. Dashed lines represent strength of attraction. **(I-L)** Image at 40 hours of the mathematical simulation of **(I)** no attraction/simNo (N-N and A-N=0), **(J)** homophilic attraction/simHomo (N-N: 1.5, A-N: 0) – yellow arrows label IN chains, heterophilic attraction/simHetero (N-N: 0, A-N: 4), and **(L)** the mixed model/simMix (N-N: 1.5, A-N: 4). Astrocyte in magenta, IN in green. **(M)** Plot showing maximum cluster size over time of 40h in (left) simulations and (right) experiments. **(N)** Plot showing normalized number of clusters over time in (left) simulations and (right) experiments. 1 indicates 0 clusters, values close to 0 indicate many clusters. **(O)** Plot showing proportion of interneurons in clusters over time in (left) simulations and (right) experiments. 1 indicates all interneurons are in clusters, 0 indicates no interneurons are in clusters. **(P)** Immunostaining of an 8-month-old explant, 1 month after plating at the leading edge with strong resemblance to the mixed model (l). For **(M-O)** N=6 per simulation and N=6 per experimental condition.

We next investigated whether these astrocyte-IN interactions play a role in stream organization in long-term explants. Immunostaining of thin streams showed numerous GFAP+ astrocytic processes closely associating with INs (Fig. 4E). Live imaging of higher density streams revealed non-GFP+ cells lining the migrating INs, closely resembling the 3D glial tube (Fig. 4F). Consistently, electron microscopy of 7-month whole organoids identified an astrocyte encapsulating a migrating neuron in close proximity to a stream (Fig. 4G). However, the exact nature of astrocyte contribution is difficult to identify as stream formation unfolds over an extended time period, and experimental astrocyte ablation might affect intrinsic IN physiology (50). To identify the relative contributions of neuron-neuron (N-N) and astrocyte-neuron (A-N) interactions leading to stream formation, we developed a two-agent (N, A) mathematical model to simulate the emergence of migration dynamics in late explants. The model incorporated experimentally defined parameters, including number of cells, soma radius (N=10 *µ*m, A=30 *µ*m) and migration speed (see Methods). Altering N-N and A-N attraction strengths enabled us to systematically interrogate how local interactions drive large-scale organization (Fig. 4H).

We first tested whether the minimal model could reproduce experimentally observed migration dynamics. Simulations of INs without astrocytes or intercellular interactions represented the ‘Early’ condition, whereas introducing astrocytes with N–N and A–N interactions represented the ‘Late’ condition (fig. S5D). As experimentally measured migration speeds were used to calibrate the model, we assessed simulated track curvature using the L2 penalty previously analyzed experimentally (fig. S5E). Notably, introducing N–N and A–N interactions reduced the L2 penalty of INs, generating straighter simulated trajectories that recapitulated late explant experiments (Fig. 3H). Thus, despite incorporating minimal experimental parameters, the model reproduced the emergence of more directed migration through local N–N and A–N interactions.

To determine the minimal interactions required to reproduce the clusters and connected IN chains observed at late timepoints, we generated four ‘Late’ model variants with different combinations of N-N and A-N attraction (Fig. 4I-L). Astrocytes were included in all variants, with their attraction strength kept at A-A=1 arbitrary unit (a.u.). This reflected the interconnected networks astrocytes form through gap junctions(51, 52), which were observed in our organoids (fig. S3F). We tested: No Attraction (N-N=0; A-N=0; simNo), Homophilic-only (N-N=1.5, A-N= 0 a.u.; SimHomo), Heterophilic-only (N-N=0, A-N=4 a.u.; SimHetero), and Mixed-attraction conditions (N-N=1.5, A-N=4; SimMix) (Fig. 4I-L, Supplementary Videos 8-10). SimNo and simHetero produced dispersed neurons resembling early migration, while N-N attraction in simHomo resulted in chain formation. The combination of N-N and A-N in simMix generated both chains and interconnected cellular clusters.

Each model was benchmarked against the experimental Early (expEarly) and Late (expLate) time-lapses using three quantified parameters. First, the models were analyzed for the maximum cluster size over simulation time (Fig. 4M). While simNo and simHomo maintained low cluster sizes, simMix and simHetero both reached maximum cluster sizes comparable to ExpLate over 40 hours. This suggests large cluster sizes can be achieved through A-N interactions alone. However, only simMix recapitulated both the normalized number of clusters (Fig. 4N; Methods) and the proportion of INs within clusters observed in expLate experimentally (Fig. 4O). Therefore, our model indicates that both N-N and A-N attraction are required to recapitulate experimental Late explant dynamics that lead to stream formation over time. Importantly, this only occurred when A-N attraction was greater than N-N, as simulations with equal or stronger N-N attraction exhibited dispersed neurons and no cellular clusters (fig. S5F,G). Thus, the simulations support a model where a weak or transient homophilic attraction between INs, potentially mediated by PSA-NCAM to allow migration(53), is coupled with a stronger heterophilic attraction with astrocytes for the emergence of the observed late migratory dynamics. Together, our explant data and mathematical model support an intrinsic propensity for late-born INs to chain migrate, while interactions with astrocytes contribute to their structural organization into streams.

### Single cell spatial and time course analysis identifies a distinct stream interneuron state

We next asked whether chain migrating INs were associated with a temporally distinct transcriptional cell state. Single-cell spatial transcriptomics (sc-spatial RNA-seq) using the Stereo-seq platform(54) allowed us to spatially identify streams to extract their transcriptional signature from 9-month CGE-Os (Fig. 5A, fig. S6). We then integrated scRNA-seq data (10X Genomics) from a time course of 2-, 5-, and 9-month-old CGE-Os, sampling the periods before and after stream formation, identifying clusters of neural progenitors, astrocytes, and oligodendrocytes, in addition to immature and mature inhibitory neuronal lineages (fig. S7A-E). A stream module score from the spatial data was then computed for the 9-month-organoid to identify the equivalent stream population from scRNA-seq clusters (Fig. 5B,C, fig. S7F-J).

**Fig. 5.**
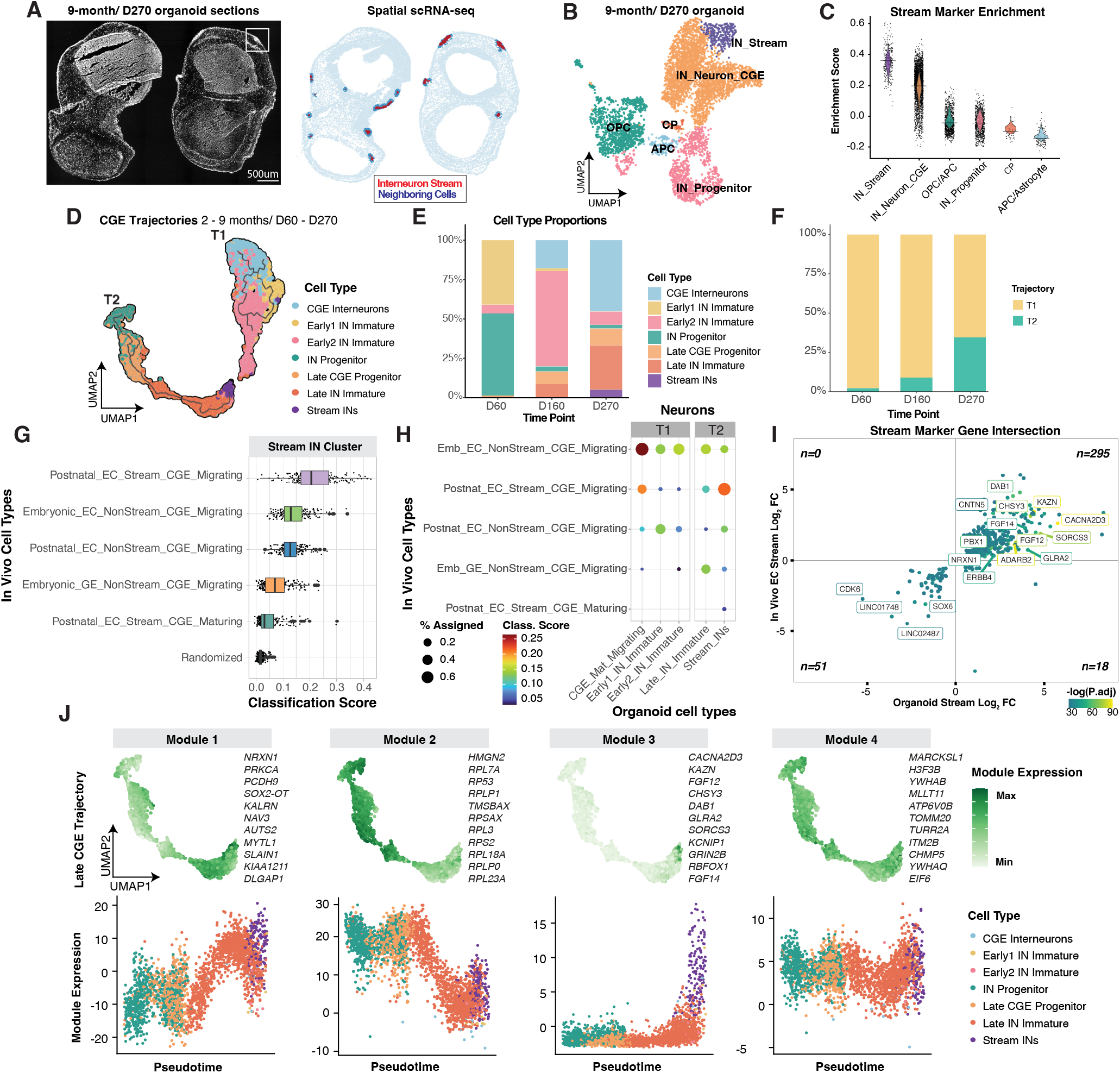
Single cell spatial and time course analysis identifies a distinct stream IN state. **(A)** Left: Spatial single-cell transcriptomic profiling of 9-month-old (9 m.o.) CGE-enriched organoid sections. Right: Single-cell spatial embedding, with streams labeled in red and stream neighbors in blue. **(B)** scRNA-seq (10x) UMAP embedding of 9 m.o. CGE-enriched organoids. **(C)** Stream marker enrichments (module score) in 9 m.o. organoids by cluster. **(D)** Integrated CGE trajectories (Monocle3) of 2, 5, and 9 m.o. organoids. **(E)** Cell type composition of the integrated CGE lineages across the 3 timepoints. **(F)** Contribution of neurons from each timepoint for trajectories T1 and T2. **(G)** Classification scoring of the organoid stream from singleCellNet training on the human EC Stream (Nascimento et al., 2024). **(H)** Proportion and classification score (confidence) of singleCellNet assignments for the neuronal CGE clusters in organoids. **(I)** Intersection of *in vivo* EC Stream and organoid stream markers significant in both datasets. Point color indicates mean FDR-corrected p-value. Quadrant totals are indicated. **(J)** Top: Module expression within the late CGE partition (T2), with hub genes listed. Bottom: The expression (ME) of four modules across pseudo-time in the late CGE partition.

As stream INs are late-born and exhibit distinct characteristics compared to early-migrating CGE INs, we asked whether they emerge from a later developmental trajectory. Monocle3 analysis of CGE lineage cells across the three time points identified two separate pseudo-time trajectories (Fig. 5D, fig. S8A-F). T1 comprised immature CGE INs from D60 and D160 CGE-Os terminating with maturing and migrating, but non-streaming neurons from D270 CGE-Os. T2 comprised IN progenitors and immature CGE INs from D160 and D270 CGE-Os, terminating with the IN stream (Fig 5D-F, fig. S8A-F). These analyses place stream INs along a distinct late developmental trajectory.

We next performed differential expression (DE) analysis comparing cells in the late migratory stream to all other immature or migrating CGE INs across timepoints (fig. S8G). GO enrichment analysis revealed significant enrichments for neuronal projection and synapse-related terms, indicating a potential role for adhesion and neuronal maturity-related genes in the stream, despite their immature, migratory state (fig. S8H).

The recently identified human-specific EC stream was described to deliver immature CGE INs to the entorhinal cortex until 11 months after birth(5). Late-migrating organoid streams closely recapitulate many of the defining features of the EC stream, with late-emerging chains of migrating, newborn SCGN+/COUP-TFII+ INs surrounded by a glial tube. We therefore investigated whether our stream cluster from the combined sc-spatial RNA-seq and sc-RNA-seq dataset shared transcriptional features with the *in vivo* EC Stream. We utilized a single-nucleus RNA-seq dataset that included the dissected early postnatal EC stream, embryonic GEs, and entorhinal cortex from late embryonic and postnatal developmental timepoints(5). A random forest classifier, Single-CellNet, which was trained on timepoint-resolved cell type labels from these data, classified the organoid stream as most similar to the postnatal EC Stream, with little similarity to other *in vivo* cell types, including other migrating, but non-streaming CGE INs (Fig. 5G,H, fig. S9A-C). Conversely, non-streaming organoid CGE INs in trajectory T1 were most similar to embryonic, non-streaming migratory CGE neurons (Fig. 5H). These results demonstrate the transcriptional similarity and specificity of the organoid streaming state with the postnatal streaming state as compared to other CGE migratory states.

We next performed DE using the *in vivo* dataset, by comparing the streaming cluster to other migrating CGE INs, mirroring our earlier approach in organoids (see Methods). Upon intersecting these *in vivo* markers with the organoid markers, we found that, for DEGs significant in at least one dataset, 1574 DEGs (78.7%) were directionally concordant (fig. S9D). When only the genes significant in both datasets were considered, a stronger correspondence was observed, with 295 shared positive and 51 shared negative markers, whereas only 18 genes were directionally discordant (i.e. 95.05% correspondence; Fig 5I). These analyses demonstrate a shared transcriptional state between organoid migratory streams and perinatal CGE IN chain migration in the human brain.

We next sought to identify the gene networks that define organoid chain migration. Weighted gene network correlation analysis (WGCNA) revealed 6 non-overlapping modules of co-expressed genes in the CGE lineages enriched for various gene ontologies (GO), including neuron migration, peptide biosynthesis, and ion channel activity (Fig. 5J, fig. S9E,F). We found that Module 1 and Module 3 were elevated later in pseudo-time, peaking in the organoid stream cluster (Fig 5J). While Late CGE Module 1 was broadly associated with general neuronal migration, increasing as progenitors differentiated into newborn neurons, Module 3 expression was selectively enriched in the streaming population, identifying a transcriptional network associated with the streaming state.

### Perturbation of stream module regulators disrupts distinct features of chain migration

Since Module 3 is a co-expression network upregulated in streaming INs, we reasoned that its key regulators may modulate IN chain migration. Network hubs were identified by the highest-ranked module eigengene-based connectivity (kME) (Fig. 6A). We employed Gene Set Enrichment Analysis (GSEA) on all module-associated genes ranked by their correlation with Module 3 kME, testing for specific enrichments in GO terms and NDD associations (fig. S10A). Module 3 was significantly enriched for genes associated with ASD risk and epilepsy (Fig. 6B). To test if these findings were preserved in the *in vivo* EC stream, genes within Module 3 were intersected with the previously identified EC Stream markers. This revealed that the majority of Module 3 genes were preserved as positive markers for the EC stream, including several ASD-associated genes (Fig. 6C). This indicates that the organoid stream network hubs are conserved *in vivo* and suggests a potential role for ASD-associated genes in the chain-migratory cell state.

**Fig. 6.**
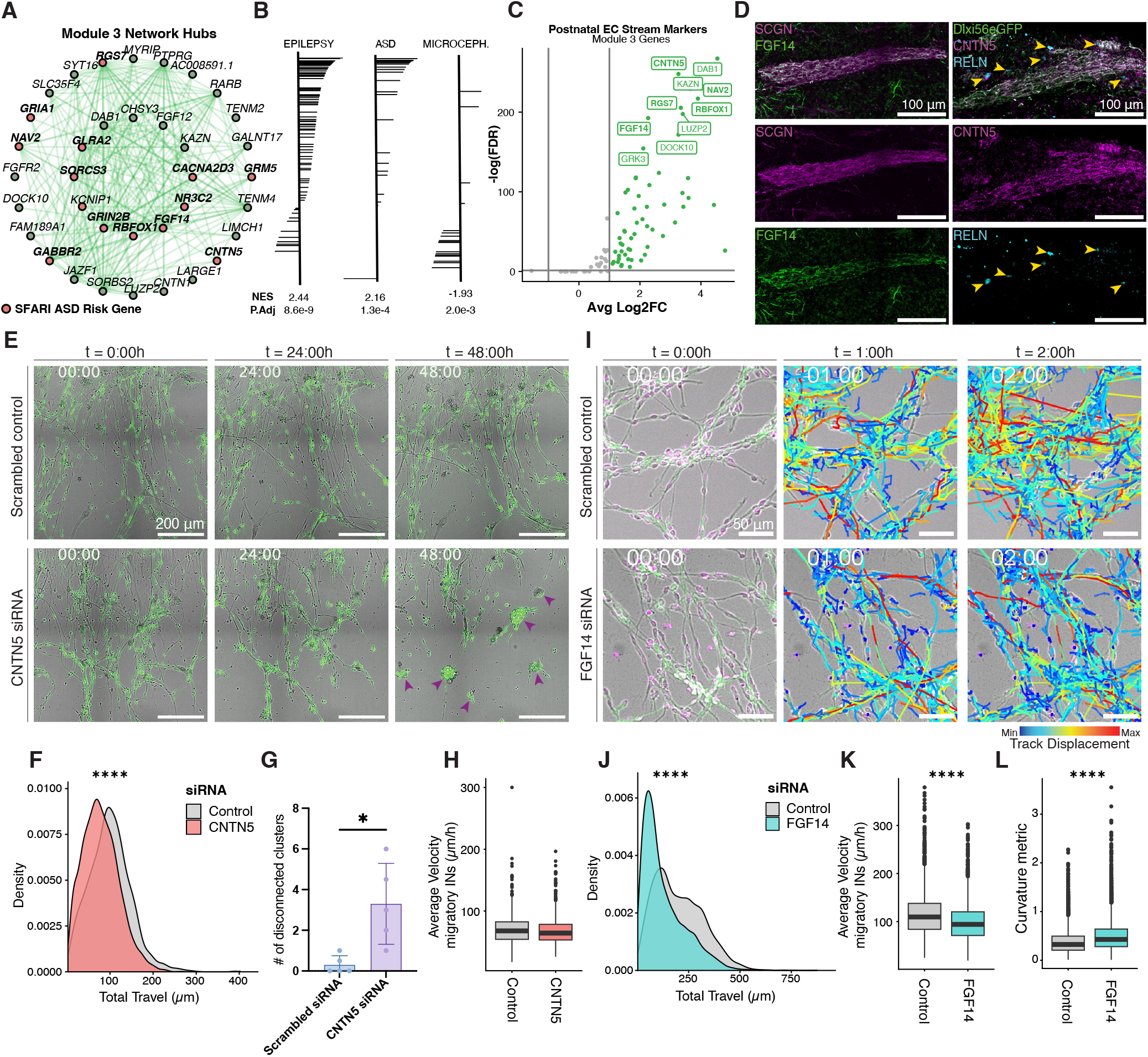
Perturbation of stream module regulators disrupts distinct features of chain migration. **(A)** Network plot of highly connected hub genes within Module 3. Red nodes mark SFARI ASD risk genes. **(B)** Gene set enrichment of NDD-associated genes ranked by correlation with Module 3 expression (kME). Positive associations with epilepsy and ASD, and negative associations with microcephaly. **(C)** Intersection of Module 3 genes with human EC Stream marker differential expression. ASD-risk genes are bolded. **(D)** Immunostainings of organoid streams for (left) *FGF14* and SCGN, and (right) *CNTN5* and RELN in 9-month and 6-month organoids, respectively. **(E)** 0-, 24- and 48h stills from time-lapse imaging video after 10-month explant KD with (top) scrambled control and (bottom) *CNTN5* siRNA. Purple arrows mark disconnected clusters. Density plot showing total travel of all IN tracks comparing control and *CNTN5* KD. *CNTN5* KD results in a significant reduction in total travel. p = 5.52×10-32. **(G)** Bar plot showing number of disconnected clusters after *CNTN5* KD. N=5. **(H)** Box plot showing average velocity of highly migratory tracks comparing control and *CNTN5* KD. Average velocity is not significantly different after filtering at p = 0.999. **(I)** 0-, 1- and 2h stills from time-lapse imaging video after 10-month explant KD with (top) scrambled control and (bottom) *FGF14* siRNA. Tracks are colored by displacement, with blue to red marking short to long tracks. **(J)** Density plot showing total travel of all IN tracks comparing control and *FGF14* KD. *FGF14* KD results in a significant reduction in total travel. p = 3.71×10-63. **(K)** Box plot showing average velocity of highly migratory tracks comparing control and *FGF14* KD. Average velocity is significantly different even after filtering at p = 5.49×10-65. **(L)** Box plot showing curvature metric/ L2 penalty of highly migratory tracks comparing control and *FGF14* KD. *FGF14* tracks are significantly more curved after filtering at p = 2.40×10-59. (F, H) N=6 for control, N=8 for *CNTN5* siRNA. **(J-L)** N=6 for control, N=8 for *FGF14* siRNA.

From these preserved network hubs, we investigated Contactin-5 (*CNTN5*) and Fibroblast Growth Factor 14 (*FGF14*), both previously uncharacterized in the context of IN migration. Immunostaining confirmed protein expression of both *CNTN5* and *FGF14* in IN streams in late organoids (Fig. 6D, fig. S10B). We therefore functionally assessed the roles of *CNTN5* and *FGF14* on migration via RNAi in late organoid explants, analyzing the effects with time-lapse imaging. Transient siRNA knockdown (KD) of *CNTN5* disrupted chain network organization, with INs aggregating into multiple disconnected, non-migratory clusters (Fig. 6E-G). Conversely, RNAi controls yielded neuronal clusters that remained interconnected in a dynamic network. Quantification of 2-minute interval time-lapses after *CNTN5* KD revealed that fewer neurons migrated compared to control, with 26.5% of INs migrating less than 50 *µ*m compared to 14.6% in control (Fig. 6F, fig. S10C). When non-migratory INs within disconnected clusters were excluded, average velocity and track curvature were not significantly altered, suggesting that the phenotype occurs through changes in structural organization rather than migratory dynamics (Fig. 6H, fig. S10D).

In contrast, KD of *FGF14* markedly reduced IN migration, with 24.3% travelling less than 50 *µ*m compared to 7% in controls but did not result in observable changes in chainnetwork architecture (Fig. 6I, fig. S10E,F). Among migrating INs, there was a significant decrease in average velocity of 13.5% and a significant increase in track curvature (Fig. 6K,L). Following each time-lapse experiment, KD was confirmed with paired immunofluorescence analysis to observe reductions in protein levels (fig. S10G,H). Using this system, we thus identified distinct functions for two stream markers, where *CNTN5* regulates chain network organization and *FGF14* regulates migratory dynamics. As these genes are conserved in the EC stream, our data suggests they may play similar roles in postnatal IN migratory streams in human development.

## Discussion

Snapshots into the neonatal human brain have revealed multiple migratory streams targeting the prefrontal, cingulate, and temporal cortices(2, 3, 5, 55), areas strongly involved in advanced functions encompassing cognition, emotion, and memory(56–58). Notably, these areas undergo a prolonged developmental period(59, 60) that coincides with late IN arrival. This raises the possibility that postnatal migratory streams contribute to the maturation of higher-order cortical systems, emphasizing the need for mechanistic characterizations in a human-specific model. Cerebral organoids have been shown to replicate human development into early postnatal stages(17, 21, 23, 24, 41), although few studies have explored postnatal structural phenomena. Here, we described the formation of a late-born CGE IN stream that arises at 6 months and persists at 17 months in cerebral organoids. This provides a platform to investigate the live migration dynamics of a unique developmental process occurring during a critical perinatal window.

Our model reveals that stream formation emerges through intrinsic properties of late-born INs, coupled with astrocyte interactions. *In vivo*, the postnatal SVZ provides a multicellular neurogenic niche from which streams arise, with ciliated ependymal cells lining the ventricles and regulating adult neurogenesis and migration(5, 61), while blood vessels provide a migratory scaffold and chemogenic cues(3, 49). However, streams robustly formed in organoids in the absence of vasculature and ependymal cells, as confirmed by electron microscopy and transcriptomically, indicating they are not necessary for stream formation itself. Furthermore, streams exhibited chain migration independently of a dorsal component in both whole CGE-Os and in explants removed from the 3D niche. This highlighted the importance of cell lineage, as birth-dating and time-course analyses show that streams emerge from a distinct late-born CGE population, which has the intrinsic capacity for chain migration. This subpopulation had distinct migratory properties, including higher velocity compared to early-migrating INs. In the postnatal human brain, this may allow INs to travel extended distances to reach their final destinations(46).

Astrocytes, however, were shown to contribute to the organization of IN streams. *In vivo*, astrocytes provide an important structural scaffold and chemogenic cues to postnatal streams(33), which was structurally recapitulated in organoids and explants. Importantly, late INs formed chains before encountering astrocytes, whereas astrocytes subsequently associated with INs in dense clusters and lined 2D streams following long-term plating. Consistently, mathematical modelling showed that N-N attraction alone was sufficient for chain formation, whereas reproducing the full phenotype with cluster formation required stronger A-N attraction as well. By recapitulating the temporal progression of human neurodevelopment *in vitro*, organoids revealed the sequence and minimal cellular interactions underlying stream formation, from the intrinsic emergence of IN chains to the larger organization with astrocyte interactions.

Our live system further allowed stream manipulation by transient perturbation of stream-specific regulators. Differences across phenotypes following KD of *CNTN5* and *FGF14* indicate that the mechanisms governing migration dynamics can be uncoupled from structural organization. *FGF14* intracellularly acts as a voltage-gated sodium channel modulator(62) and is notably required for adult neurogenesis and maturation of new neurons in the dentate gyrus (DG)(63). We observed that *FGF14* KD reduces late-IN migration velocity without affecting overall architecture, suggesting a role in regulating intrinsic migration dynamics. Whether *FGF14* acts through a common mechanism to regulate adult neurogenesis in the DG and the V-SVZ remains to be determined. In contrast, the architectural changes from *CNTN5* KD, a neural cell adhesion molecule, suggests a broader role in intercellular organization. As neither gene has previously been implicated in cortical IN migration, it is possible these roles are particularly relevant to late chain migration.

The substantial overlap between stream-modulating genes and ASD-associated genes suggests that this late developmental process may be additionally susceptible to NDD-related perturbations. Both *CNTN5* and *FGF14* are ASD-associated(64–66), with *CNTN5* implicated through neurite outgrowth and synaptogenesis(67, 68), and *FGF14* as an intracellular regulator of neuronal excitability and synaptic plasticity(69, 70). While ASD has traditionally been viewed as a circuit-level disorder (71–73), our results demonstrate that these ASD-associated genes can regulate late-IN migration as well, potentially extending the list of neurodevelopmental processes implicated. Furthermore, *FGF14* SNP mutations have been associated with changes in EC- and hippocampal volume in Alzheimer’s disease (AD) patients(74, 75). This is particularly relevant since the EC, one of the earliest affected brain regions in AD(76), receives immature INs via the EC stream. Nascimento and colleagues have suggested a selective vulnerability of INs from the EC stream in AD, while *FGF14* may potentially exert neuroprotective effects(77), which would be interesting to investigate further. Our *in vitro* stream model opens new avenues for investigation. While our CGE-O patterning approach yielded streams that recapitulate the CGE-enriched EC stream, by expanding the protocols to include different cell types, we may model the diversity of MGE- and LGE-derived INs within the Arc(55) and MMS(2). This would enable comparative analyses across distinct perinatal IN streams, allows us to distinguish the mechanisms intrinsic to chain migration from those that are lineage specific. The astrocyte lineage would be another interesting direction to explore. In the postnatal rodent brain, the rostral migratory stream (RMS) is a well characterized example where migratory neuroblasts (type-A cells) arise from astrocyte-like adult neural stem cells (type-B cells)(78), thereby sharing a common lineage. Whether a similar lineage relationship exists in human postnatal migratory streams is unknown. Notably, the RMS is composed of a separate IN population from that of our organoid stream that is COUP-TFII negative(7). Nevertheless, the structural similarities raise the possibility of shared developmental trajectories that could be investigated with lineage tracing. In the context of NDDs, we have analyzed two genes but identified many others that are implicated in ASD and epilepsy. Our system allows their functional examination on late-migration and on broader effects in late brain development, as well as exploring personalized treatment options such as gene therapy.

In the wider context of human brain evolution, our system enables functional investigation of late-born INs beyond migration. A key question is how late-born CGE INs specifically shape cortical networks. Network plasticity is thought to require GABAergic input for experience-dependent circuit refinement(79, 80). While reports on IN involvement during postnatal plasticity periods largely focus on MGE-derived Parvalbumin INs(81), CGE-derived INs may play a more nuanced role. Although transplantation of mouse immature CGE-derived neurons did not induce postnatal cortical plasticity(82), this may differ in humans, where CGE INs are proportionally expanded and may exert broader functions(6, 7). Indeed, VIP+ IN-mediated disinhibition actively facilitated plasticity during critical learning periods(83, 84), highlighting their potential significance. With our established model, the functional effects of late-born IN integration into developing neuronal circuits can be studied through late assembloids and animal transplantations to assess for changes in activity.

Our findings demonstrate that cerebral organoids can recapitulate human neurodevelopmental processes occurring into early postnatal stages. By capturing the temporal emergence and progressive organization of migratory streams, organoids enabled us to reconstruct their developmental sequence and mechanistically dissect a process previously inaccessible in the human brain. These results position organoids as a platform to interrogate late developmental processes in humans, which may underlie evolutionary cognitive expansions while simultaneously increasing neurodevelopmental vulnerability.

## Supporting information

Supplementary Video 1. Chain migration in organoid streams

Supplementary Video 2. Early organoid explant migration

Supplementary Video 3: Late organoid explant migration

Supplementary Video 4: Early organoid explant tracking

Supplementary Video 5: Late organoid explant tracking

Supplementary Video 6: Interneurons interact with astrocytes in late explants

Supplementary Video 7: Interneurons migrate in chains without astrocytes in late explants

Supplementary Video 8: Homophilic model N-N=1.5

Supplementary Video 9: Heterophilic model A-N=4

Supplementary Video 10: Mixed model N-N=1.5, A-N=4

## Acknowledgments

We thank A. Piszczek (VBCF Histology Facility) and L. Pu (BGI Research) for assistance with sample processing for STOMICs spatial transcriptomics experiments, and the VBCF Histology Facility team. We thank the BioOptics Core Facility, specifically P. Pasierbek, A. M. Cencerrado, G. Bradamante, and T. Lendl for assistance with image acquisition and processing; the Next Generation Sequencing Facility at ViennaBioCenter Core Facilities (VBCF) for single-cell sequencing services; the IMBA/IMP/GMI Bioinformatics for sequencing analysis. We thank D. Reumann for assistance with 3D printing of PDMS moulds. We thank C. Li for assistance with 10x single cell transcriptomics sample preparation. We thank M. Paredes for insightful discussions.

## Author Contributions

Conceptualization: SNW, NSC, JAK

Methodology: SNW, MF, LM, LBM, SG-G

Formal analysis: SNW, MAN, MF, LM

Investigation: SNW, MAN, MF, LM, MN, EN, DvdH, LBM, SG-G

Data curation: MN, MAN, MF, LM

Visualization: SNW, MAN

Funding acquisition: JAK

Supervision: NSC, JMG-V, SM-A, JAK

Writing – original draft: SNW

Writing – review & editing: SNW, MAN, NSC, JAK, with input from all authors

## Materials and Methods

Biorender was used for generation of schematics in figures.

### Patient sample selection and cell line generation

Feeder-free hESC line WA09 (H9 ESCs) was commercially obtained from WiCell. HiPSC line 1 is an iPSC line reprogrammed in-house from fibroblasts of healthy donors (internal nomenclature: iPSCs 176/1). hiPSC lines 2 and 3 were generated in-house and published in Eichmüller et al. 2022. The study was approved by the local ethics committee of the Medical University of Vienna (MUV). The Dlxi56-eGFP en-hancer was inserted into the AAVS1 locus of all cell lines as previously described(25).

### Culture of pluripotent stem cells

PSC culture was performed as published previously, with minor modifications(25). Feeder-free human ESCs and iPSCs were cultured in 6-well plates (Corning, cat. 3516) coated with hESC-qualified Matrigel (Corning, cat. #354277). Media used was Essential8 (E8) medium (Thermo Fisher, cat. #A1517001) supplemented with BSA. Cells, when 70% confluent, were passaged by incubation in 1 mM EDTA (Pan-Reac AppliChem, cat. #A1103) in sterile D-PBS without calcium and magnesium (Thermo Fisher, cat. #14190136) for 4 min at 37°C and subsequent passaged to a Matrigel-coated plate with E8 medium. Cells were routinely checked for mycoplasma and karyotyped.

### Organoid generation: Extended Ventral (MGE-enriched), Short ventral (CGE-enriched), and Dorsal patterning

Briefly, cells at 70% confluency were treated with 1 *×* Accutase solution (Sigma-Aldrich, cat. #A6964) for 4 min at 37°C and resuspended in stem cell medium: E8 medium supplemented with Rock Inhibitor Y-27632 (Stemcell Technologies) for cell counting. For dorsal, short ventral, and extended ventral embryoid body (EB) generation, 9,000 cells were added to each well of a 96-well low-attachment U-bottom plate (Corning, cat. #COR7007) in 150 *µ*l of stem cell medium. On day 3, dorsal forebrain EBs (dorsal EBs) received stem cell medium and ventral forebrain EBs (ventral EBs) received neural induction medium (NI medium(85)). From day 5–11, dorsal EBs were cultured in NI medium only, while ventral EBs were patterned with NI medium with 100 nM SAG (Merck-Millipore, cat. #US1566660) and 2.5 *µ*M IWP2 (Sigma-Aldrich, cat. #IO536). EBs were transferred to Matrigel-supplemented media (Corning, Cat. #354234) on day 11 in 10 cm dishes (TPP, cat. #Z707651). On day 13, medium was changed to either improved differentiation medium-A supplemented with 1% Matrigel (Imp-A + MG) and 3 *µ*M CHIR99021 (Merck, cat. #361571) for dorsal EBs, into Imp-A + MG, 100 nM SAG and 2.5 *µ*M IWP2 for extended ventral, or only Imp-A + MG for short ventral protocol. Dorsal organoids received an additional CHIR99021 (3 *µ*M) boost on Day 15, while extended ventral received IWP2+SAG until Day 17. All organoids received IMP+A on day 20 and media was changed every 3-4 days for long term culture.

### Day 120 organoid fusion

Short ventral and dorsal organoids were fused together at day 120 to analyze directed late migration. PDMS molds were sterilized overnight in 70% ethanol (EtOH), then washed twice in 1xPBS to remove all traces. Mold indents where organoids will be placed were coated with anti-adherence rinsing solution (Stemcell Technologies, Catalog #07010), then washed 2x with 1xPBS. Organoids were placed into the mold indents, keeping track of the dorsal and ventral orientation i.e. dorsal organoids placed on the left and ventral on the right. Media was removed with a P200 to ensure organoids were pressed together with no media remaining in the indent. Organoids were covered in a droplet of Matrigel and incubated for 20 mins at 37°C, 5% CO2. After Matrigel solidification, fused organoids were carefully flushed out with a P1000 into fresh media. Fused organoids are very delicate at this stage and are left non-shaking for one week and fed twice a week, before placing on shaker at 57 rpm.

### Birth dating, EdU addition

A working concentration of 10 *µ*M was used. EdU was diluted from stock concentrations directly into organoid media and organoids were incubated for 2 hours. Organoids were then washed 2x with 1xPBS and placed in fresh media. Organoids were fixed 10 days later to image EdU+ cells. Imaging was performed following the protocol for the Click-iT EdU Cell proliferation Kit (Invitrogen, C10340).

### Immunohistochemistry on organoids

Organoids were fixed in 4% paraformaldehyde (PFA) for 1 hour at room temperature (RT), then washed 3x with 1xPBS and equilibrated in 30% sucrose in PBS at 4°C overnight. Organoids were embedded in OCT (optimal cutting temperature compound; Scigen, Cat. No. 4586) and sectioned at 20 *µ*m using a cryostat (Leica NX70). Sections were dried overnight at RT then stored at -20°C. To stain, sections were thawed to RT, then blocked and permeabilized in blocking solution (0.25% TritonX-100 with 4% normal donkey serum) (Millipore) in PBS for 1 hour at RT. Sections were then incubated overnight at 4°C with primary antibodies in staining solution (0.1% Triton X-100 with 4% normal donkey serum) (Millipore) in PBS. Sections were washed 3x in 1xPBS. Secondary antibodies were incubated for 2h at RT and afterwards sections were washed 3x in 1x PBS. DAPI was added to secondary antibodies to mark nuclei. Secondary antibodies labeled with Alexa fluor 488, 568, or 647 (Invitrogen) were used for detection. Slides were mounted using fluorescence mounting medium (Dako) and dried overnight before imaging.

### Microscopy

Widefield images were acquired using the Zeiss AxioVert A1 Inverted Microscope. Confocal images were acquired on LSM800 Axio Observer Z1 confocal laser scanning microscopes (Zeiss). Large overview images were acquired using the Panoramic Slide Scanner 250 Flash II or III system (3DHistech). Time-lapse recordings on whole organoids were performed with the Olympus Spinning Disk system based on the Olympus IX3 Series (IX83) inverted microscope with a dual-camera Yokogawa W1 spinning disk. A 40x/0.75 (Air) WD 0.5mm objective was used. Time-lapse recordings on organoids explants were performed with the Celldiscoverer 7 with Axiocam 506 -CCD (Pixel size: 4.54*µ*m) and Hamamatsu Orca Flash 4.0 (Pixel Size: 6.5 *µ*m) camera and a Plan-Apochromat 20x/0,7 objective.

### Antibodies for immunohistochemistry

COUP-TFII (PP-H7147-00; Clone H7147; R&D; 1/200), NKX2.1 (ab133737; Abcam; 1/200), CALB2 (CR6B3; Swant; 1/1000), SCGN Rabbit (HPA006641-100UL; Sigma-Aldrich; 1/1000 2D; 1/200 3D), SCGN Mouse (MA5-31337; Invitrogen; 1/1000), GFAP (ab4674; Abcam; 1/1000), S100B (ab52642; Abcam; 1/1000), Doublecortin (ab77450; Abcam; 1/200); PSA-NCAM (MAB5324; clone2-2B; Sigma-Aldrich; 1/200); EGFR (AF231; R&D; 1/200); KI67 (14-5698-82; Invitrogen; 1/100); OLIG2 (AF2418; R&D; 1/200); TBR1 (ab31940; Abcam; 1/500); RELN (MAB5366; clone142; Sigma-Aldrich; 1/200); *CNTN5* (PA5-59486; Invitrogen; 1/200); *FGF14* (75-096; cloneN56/21; Antibodies Inc; 1/200); FOXG1 (Produced in-house), LHX6 (sc-271433; SCBT; 1/500), MAP2 (PA1-10005; Invitrogen; 1/1000), DLX2 (sc-393879; SCBT; 1/200), SOX2 (MAB2018; R&D; 1/500), Chicken-GFP (GFP-1020; Aves labs; 1/1000) Goat-GFP (AB0066-20; Sicgen; 1/1000).

### Time-lapse imaging of 3D organoids

Whole organoids were selected based on GFP+ streams close to the surface using a widefield microscope. The organoid was placed in a *µ*-Plate 24 Well for imaging (Cat.No:82426; Ibidi) and filled with 2.5 ml Imp+A media. Organoids were imaged using the spinning disk confocal microscope at 40x magnification of stream regions with multiple Z-planes to capture the stream. Images were acquired at ∼5-10% laser power to prevent phototoxicity of live neurons. Images were acquired continuously for hours/ days with imaging every 10-20 minutes.

### Organoid explant assay: plating organoids and time-lapse imaging

Organoids were placed on a dish with minimal media and dissected into ∼2mm pieces with P10 long pipette tips. Cut organoids were washed 2x in 1x PBS to remove debris and placed non-shaking in the incubator. Cut organoids were washed again the next day to remove debris and then fed twice a week. After at least 10 days of recovery, cut organoids were plated on Matrigel coated 4-well cell culture lumox chambers (Sarstedt, cat. #94.6150.401) with a droplet of Matrigel. Interneuron migration was imaged from 5 days on with the Celldiscoverer 7. Time-lapses were conducted for 3-5 days at 20x magnification with images acquired every 20 minutes. In the case of time-lapses for interneuron tracking, images were acquired every 2 minutes for 3 hours at lowest laser power to prevent phototoxicity. Before imaging, chambers were incubated with BioTracker 650 Red Nuclear Dye to identify all neurons for tracking (1:20,000; Sigma-Aldrich, cat. #SCT119).

### Immunohistochemistry on 2D explants

After live imaging, migrated neurons were directly fixed in chambers with 4% PFA for 1h at RT, then washed 3x with 1xPBS and stored sealed at 4°C until immunostaining. To stain, 300ul permeabilization solution was added per chamber for 30 mins at RT. Chambers were then incubated with primary antibodies in staining solution overnight at 4°C, or for 2h at RT. Chambers were washed 3x with 1xPBS. Secondary antibodies were incubated for 2h at RT then washed 3x with 1xPBS. DAPI was added to secondary antibodies to mark nuclei. After washing with 1xPBS, peel off the frame and use a scalpel to cut the edges of the lumox layer containing migrated neurons and transfer the whole layer onto a glass slide. Mount cover glass on top with mounting medium and dry at RT overnight before imaging.

### Quantification of 2D migration tracking

Cell trajectory analysis was performed using the TrackMate(86, 87) plugin within Fiji (ImageJ), with subsequent detailed filtering and quantitative analysis conducted in Wolfram Mathematica. To ensure robustness of the results and focus the study on actively migrating neurons, a multi-step filtering process was implemented. The filtering criteria included a minimum threshold for the number of spots per track to ensure that analyzed cells were observable for a significant duration of the video recordings. Furthermore, permissive thresholds for the total traveled distance and the net displacement were applied to maximize the number of tracks included in the analysis while specifically focusing on moving neurons. This configuration was chosen to maintain a stable data basis for population-level characteristics without excluding datapoints (fig. S4H). For the phenomenological comparison between experimental conditions, the analysis primarily focuses on average migration speed and path straightness. A key metric for assessing the shape and quality of these trajectories is the normalized L2 penalty. This metric represents the “jitter” or “bendiness” of a track. Unlike standard total curvature, the penalty mathematically penalizes sharp, abrupt turns more heavily than gradual directional changes. Consequently, low values indicate smooth trajectories without frequent, sharp hooks or sudden adjustments as visualized in fig. S4I,J.

### RNAi of organoid explants

*CNTN5* (MCE; HY-RS02930) and *FGF14* (ThermoFisher; 4392420) siRNAs were diluted in media without antibiotics to final amount of 10pmol and added to chambers. Time-lapse imaging was performed 3 days after siRNA addition to evaluate the phenotype and fixed directly after imaging to evaluate knockdown efficiency.

### Quantification of COUP-TFII and NKX2-1 in streams

Tif files of streams were processed with Fiji (ImageJ). In Fiji, the freehand selection tool was used to draw a perimeter around Dlx5/6+ streams in the GFP channel. Streams were then isolated using ‘edit, clear outside’ to remove non-stream cells. Cell type classification was then performed using a custom macro. Briefly, nuclei segmentation was performed on the DAPI channel with Cellpose(88). Channels 2 and 3 were then overlayed on the segmented nuclei and identified as positive or negative with intensity thresholding. Automated detection was visually verified, and cell counts were normalized to total number of DAPI cells within each stream.

### Quantification of siRNA knockdown

All immunofluorescence image acquisition was performed with the same microscope settings (laser power, exposure time) between paired control and siRNA conditions. Processing and quantification was performed in Fiji (ImageJ). Briefly, tif files were split by channel, and thresholding was applied to the GFP channel to isolate individual interneurons a a binary. Then, ‘analyze particles’ was selected to count each neuron as an ROI. After mask creation, ‘set measurements’ on the siRNA fluorescence channel to extract mean fluorescence intensity per ROI. Each ROI intensity value was normalized against an average of 6 background regions per image.

### Transmission electron microscopy (TEM)

Brain organoids were fixed with 3.5% glutaraldehyde. Samples were postfixed with 2% osmium tetroxide solution. They were dehydrated in crescent ethanol concentrations, stained with 2% uranyl acetate and embedded in araldite resin (Durcupan ACM Fluka, Sigma), followed by polymerization at 70°C for 72 h. Serial semi-thin sections (1.5 *µ*m thick) were cut using an Ultracut UC-7 ultramicrotome (Leica, Heidelberg, Germany), mounted onto slides, and stained with 1% toluidine blue to identify regions of interest. Selected semi-thin sections were glued to resin blocks and detached from the slides by repeated freezing in liquid nitrogen. Ultrathin sections (70–80 nm) were then cut from the blocks with detached semithin material and collected on copper grids. Ultrathin sections were contrasted with lead citrate (Reynolds’ solution) and examined under a FEI Tecnai G2 Spirit transmission electron microscope (FEI Europe, Eindhoven, Netherlands). Images were acquired using Radius software (Version 2.1) with a XAROSA digital camera (EM-SIS GmbH, Münster, Germany). Consecutive series between 120-200 ultrathin sections were collected from migrating chains and surrounding astrocytes. This serial imaging approach enabled three-dimensional reconstruction of entire cells, including their nuclei, mitochondria, and membrane specializations, providing spatial context for ultrastructural features.

### Correlative light and electron microscopy (CLEM)

For CLEM of SCGN, GFAP, and ultrastructure, brain organoids were fixed in 3.5% glutaraldehyde, post-fixed and embedded as described above. Samples were sectioned into series of three consecutive semi-thin sections (1.5 *µ*m) on gelatin-coated slides using an ultramicrotome. Post-embedding immunohistochemistry for GFAP and SCGN labelling was performed on the first and third sections, respectively, to define regions of interest. First, the resin was removed from the sections using ethoxide for 12 minutes, followed by rehydration through a graded series of decreasing ethanol concentrations. They were then rinsed in water and PBS, treated with 1% sodium metaperiodate in H2O for 10 minutes and with 1% sodium borohydride in PB for 15 minutes to reduce glutaraldehyde-induced cross-linking. After rinsing in PBS, the sections were incubated in a blocking solution containing 10% normal goat serum (NGS) (Sigma, San Luis, MO, USA) in PBS for 30 min. Then, sections were incubated overnight at 4ºC with primary antibodies anti-GFAP rabbit (1:500, Dako) or anti-SCGN rabbit (1:1000, Sigma), respectively, diluted in the same blocking solution. The next day, sections were washed and incubated with the corresponding biotinylated secondary antibody anti-rabbit in PBS (1:200, Vector) for 50 minutes, followed by the avidin-biotin-peroxidase complex (ABC Kit, 1:200) for 45 min. DAB detection was performed with 0.05% diaminobenzidine and 0.03% H_2_O_2_ in PBS for 10 minutes. After final PBS washes, sections were dehydrated in a graded ethanol series and mounted with Eukitt. Once the region of interest was identified, selected semi-thin sections from the second slide of each series (correspondingly located between the sections labelled with GFAP and SCGN) were processed to obtained TEM images as described above.

### Single cell spatial transcriptomics sample preparation

Organoid samples were identified based on the presence of clear Dlxi56-eGFP+ streams using the Zeiss AxioVert A1 Inverted Microscope. Organoids were fixed with fresh 4% PFA in milliQ water. This was preferred over fresh-frozen methods to maintain overall organoid structure and neuronal architecture. Then, the Stereo-seq workflow(54) was followed on-site and optimized for brain organoid samples. Briefly, a representative OCT block was sectioned until the tissue was reached, then 5x 10 *µ*m sections were placed onto 5 separate F-chips. The tissue was fixed and permeabilization testing was performed on a range of time intervals to identify the optimal duration for organoid tissue permeabilization while minimizing RNA diffusion. After the optimal timepoint was determined (14 mins in organoid sections), the OCT organoid block was sectioned and each section imaged until an interneuron stream was identified with GFP under the microscope. The immediate next section was discarded due to tissue drying and the following section was placed onto the S-chip (1 cm x 1 cm) for sequencing. The placed section was then permeabilized at the previously determined time interval. Reverse transcription, cDNA purification and amplification were performed, followed by library preparation with the STOmics Library Kit and sequenced on a DNBSEQ platform.

### 10x single cell transcriptomics sample preparation

For each library, two organoids were pooled, with six organoids per timepoint. Pooled organoids were washed 2x with DPBS-/- and dissociated in 1x trypsin:accutase (1:1) solution with TURBO DNase (ThermoFisher, catalogue no. AM2238), using the GentleMACS dissociator. After dissociation, the reaction was stopped with gradual addition of DPBS-/-supplemented with 10% FBS (DPBS+10% FBS). At 4°C, samples were centrifuged for 5 mins at 400g, then supernatant removed without touching the pellet. Pellets were resuspended with 1ml of DPBS+10% FBS then filtered first through a 70 *µ*m strainer, then into FACS tubes. Cells were stained with DRAQ7 (Biostatus, DR70250) for cell viability. Live cells (Alexa 700) were sorted under low pressure (100 *µ*m nozzle) with the BD FACSAria III and collected in DPBS+10% FBS on ice. Collected cells were centrifuged at 400g for 5 mins at 4°C and concentrated to 1,000 cells/ *µ*l in ∼50 *µ*l. Sample viability was tested and only samples above 85% viability were processed. 16,000 cells per library were loaded onto a 10*×* chromium controller for targeted recovery of 10,000 cells. Libraries using the ‘Cells 10X 3’ st scRNA-seq 10X Dual Index Kit TT Set A Workflow A’ were prepared following the 10*×* user guide. Libraries were sequenced on a Illumina NovaSeq SP flow cell with demultiplexing.

### Single cell spatial transcriptomics (Stereoseq) and single cell 10x genomics analysis

10x Genomics 3’ single-cell RNA-sequencing was performed in biological triplicates, with each of three samples consisting of a 20:40:40 pool (D60:160:270) of brain organoids. Raw sc-RNAseq data were analyzed with Cell-Ranger v7.1.0 using default parameters and the GRCh38-2020-A 10x Genomics reference, extended to include the GFP sequence. Cells were demultiplexed by cell line geno-type using Souporcell (v2.4). Downstream analysis was performed in Seurat v5. Samples across timepoints were integrated using Harmony followed by UMAP embedding. STOmics libraries were processed using the STOmics Analysis Workflow (SAW) v5.5.3 by the supplier, cellbin levels were exported using Stereopy v.1.3.1, and further thresholded in Seurat, removing cells with fewer than 50 detected genes.

Cells were normalized by SCT, followed by dimensionality reduction with PCA and UMAP embedding. Cluster markers were identified via Wilcoxon Rank Sum using FindAllMarkers.

For time-resolved scRNA-seq, the stream cluster was identified using the top 200 stream markers using AddModuleScore. Clusters were annotated using FindAllMarkers and the random forest classifier singleCellNet trained on *in vivo* signatures obtained from Nascimento et al., 2023. Organoid and *in vivo* (Nascimento et al., 2023) stream markers were identified using Wilcoxon Rank Sum via FindMarkers, comparing the stream cluster to all CGE interneuron clusters, down-sampling identities to equal population sizes. Organoid and *in vivo* stream signatures were compared with single-CellNet trained on fine-labeled migratory clusters including the EC Stream from Nascimento et al., 2023. Trajectories were visualized and Pseudotime was computed using Monocle3(89) on the subset of integrated cells identified as CGE. Gene coexpression networks were identified within the CGE lineage using the hdWGCNA package(90), with soft powers optimized using TestSoftPowers. Module eigene-genes (MEs) were extracted using ModuleExprScore. Network hubs were identified by ranking kMEs using GetHubGenes. Enrichment tests were performed via EnrichR and the fgsea package.

### Mathematical model

We use an individual-based model to describe the dynamics of interneurons and astrocytes. Each cell type is modelled as a disk of fixed radius; the position of the center of the disk determines its position. Interneurons also have a direction of motion. The model takes into account various interactions and forces between cells, such as protrusion extensions for astrocytes, directional change of neurons, attraction-repulsion forces for all cells, etc. We refer the viewer to the supplementary text where a full description of the model’s components is given. All these forces are incorporated into the model, in which the dynamics of each cell are given by a stochastic differential equation complemented with jump processes. We use an Euler-Maruyama scheme to approximate the model numerically, using tools from the Python Sisyphe library(91). The size of each cell type and the speed of the interneurons have been calibrated using experimental data, while other parameters have been selected by comparing the model output to the data.

All simulations and figures related to the mathematical models are done in Python 3.15. For both simulated and experimental data, we have trajectories of interneurons with their position at specific time points. The main difference is that the simulated data allows us to have the full trajectory of all cells at each simulated time points. We use the OPTICS (Ordering Points To Identify the Clustering Structure) algorithm from the scikit.learn Python package to label each interneuron. Interneurons with the same labels (starting at 0) belong to the same clusters, while interneurons with label -1 are isolated. From this, we can compute various quantities such as proportion of neurons clustering, largest cluster size and normalized cluster number. The proportion of neurons is computed by computing the number of clustering neurons (labels different from -1) divided by total number of neurons. Normalized cluster number is the number of clusters divided by the number of cells. An isolated cell is then counted as a cluster of size 1. This means that if this number is close to 1 then there is a lot of isolated interneurons (size 1 clusters), and as it decreases the number of clusters increases.

### Statistics and reproducibility

Data for stream-related quantifications are presented as mean *±* sd. Figure legends specify N-values and P values, where each N represents an independent experiment. For differential expression and marker gene analysis, all nominal p-values were adjusted for multiple hypothesis testing correction using the false discovery rate (FDR). For RNAi experiments, migration total distance, velocity, and curvature were compared across control and KD conditions using a linear mixed-effects model with RNAi treatment as a fixed effect and replicate as a random effect.

## Code Availability

All code used in the study will be publicly available under Github: https://github.com/knoblichlab/Stream_2026 and Zenodo (mathematical modeling) https://zenodo.org/records/18740760

## Data Availability

The datasets generated during and/or analyzed during the current study are available from the corresponding author on request.

## Supplementary Figures

**Fig. S1.**
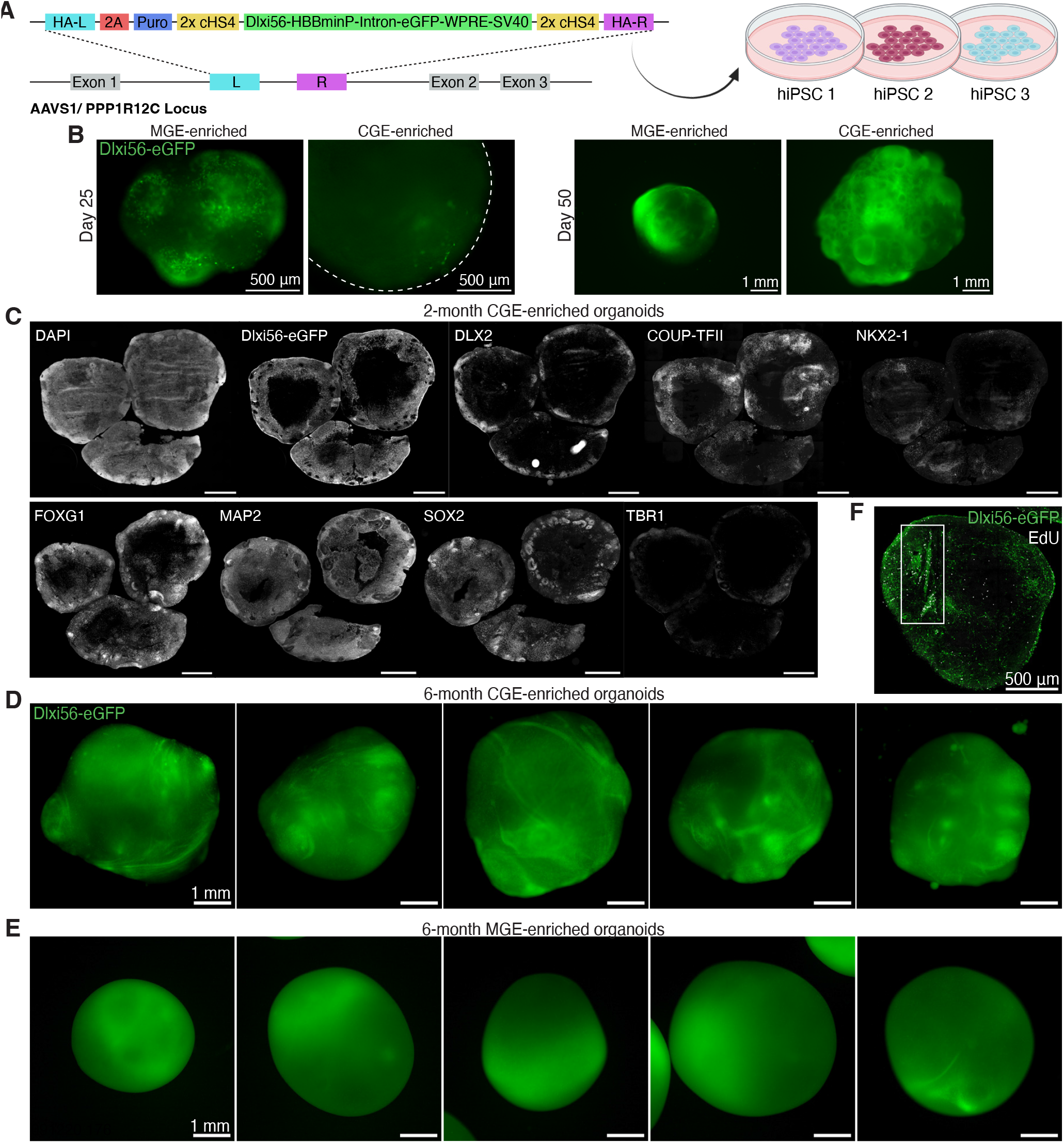
Characterization of CGE-enriched organoids. **(A)** Schematic of TALENs-based Dlxi56-eGFP reporter insertion into the AAVS1 locus in 3 hiPSC lines. **(B)** Widefield GFP images of CGE-enriched and MGE-enriched organoids at (left) day 25 and (right) day 50. Dlxi56-eGFP marks all INs. **(C)** Immunostainings to characterize whole CGE-enriched organoids at 2 months. **(D)** Widefield GFP images of CGE-enriched organoids at 6 months showing IN streams. **(E)** Widefield GFP images of MGE-enriched organoids at 6 months with minimal streams. **(F)** Immunostaining of whole organoid section from Fig. 1J of EdU+ (white) cell distribution at 7.2 months.

**Fig. S2.**
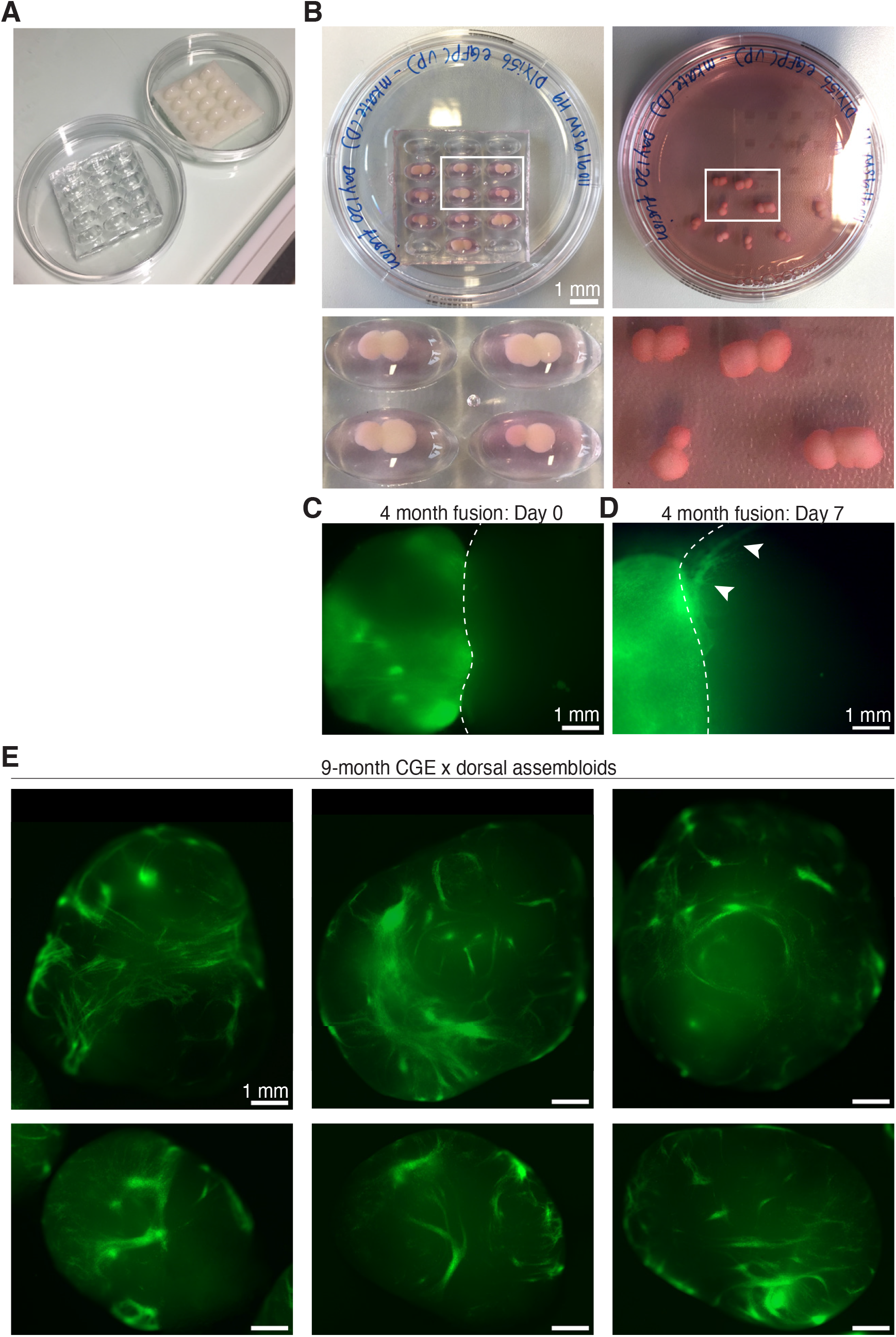
Fusion of late CGE and dorsal organoids for directed migration. **(A)** PDMS (Polydimethylsiloxane) molds were custom designed and 3D printed to allow fusion of larger organoids. **(B)** Dorsal and CGE-enriched organoids were fused by embedding in a droplet of Matrigel in PDMS molds. **(C)** Widefield GFP image of a CGE-D assembloid on the day of fusion. **(D)** Widefield GFP image 7 days post-fusion show IN streams migrating into the dorsal organoid. **(E)** Widefield GFP images of 9-month-old CGE-D assembloids.

**Fig. S3.**
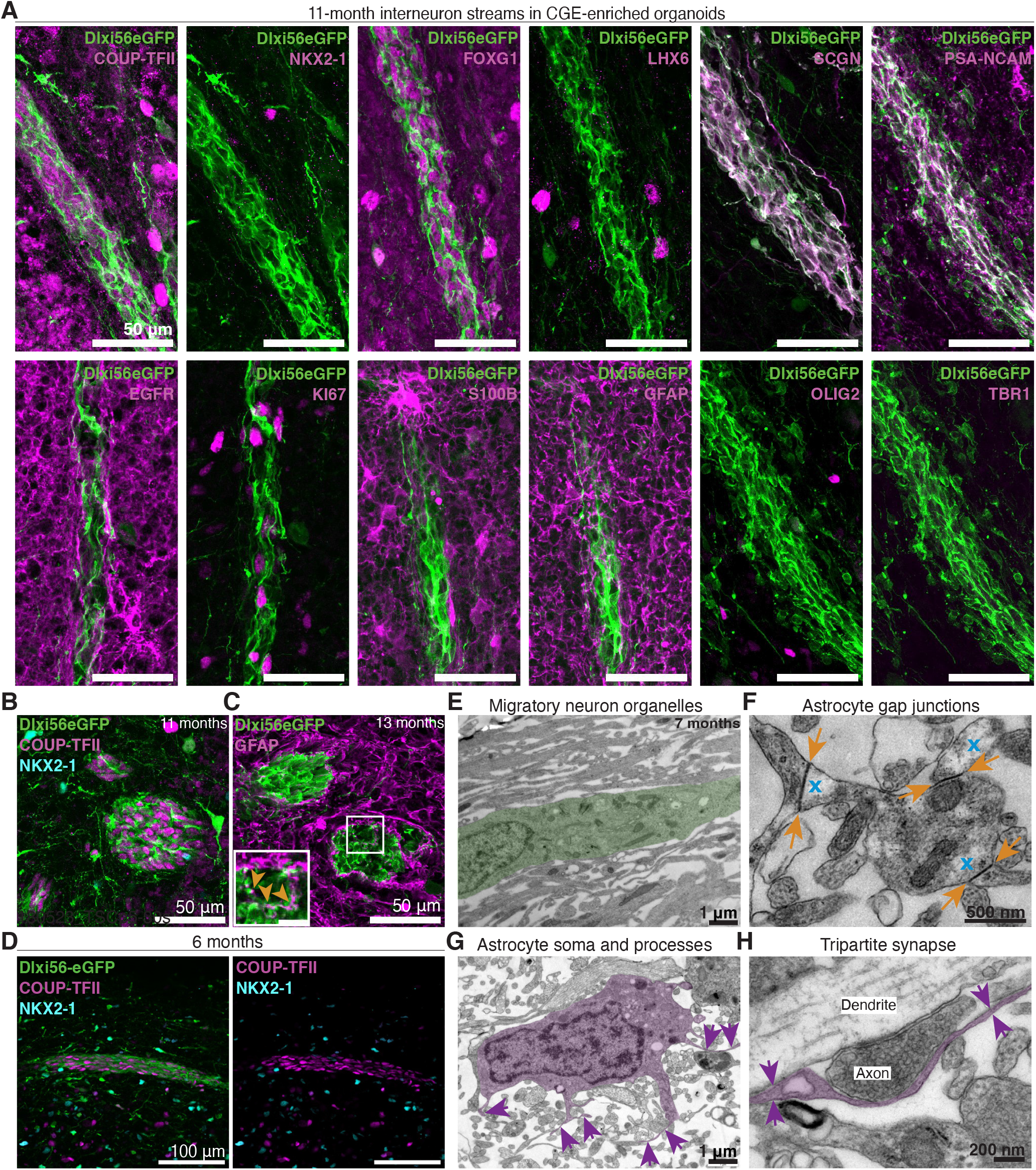
Streams are composed of CGE INs surrounded by astrocytes. **(A)** Immunostainings on 11-month organoid streams in CGE-enriched organoids. Streams are positive for COUP-TFII, FOXG1, SCGN, PSA-NCAM, and negative for NKX2-1, LHX6, OLIG2, TBR1. Streams are surrounded by S100B, GFAP, EGFR glia with few KI67 cells. **(B)** Immunostaining cross-section of stream shows COUP-TFII+ INs. **(C)** Streams are surrounded by GFAP+ glia that infiltrate processes into streams (inset – yellow arrows). **(D)** Streams in 6-month organoids are positive for COUP-TFII and negative for NKX2-1. **(E)** EM of migratory neuroblast (green) have abundant organelles like mitochondria. **(F)** Astrocytes form gap junctions in organoids visualized as dense lines (yellow arrows) and have glycogen stores, visualized as electrolucent white spaces (blue X). **(G)** Tracing from astrocyte soma (magenta) identify thin, dark processes (purple arrows). **(H)** Astrocytes form tripartite synapses in organoids (magenta, arrows) around a dendrite and axon.

**Fig. S4.**
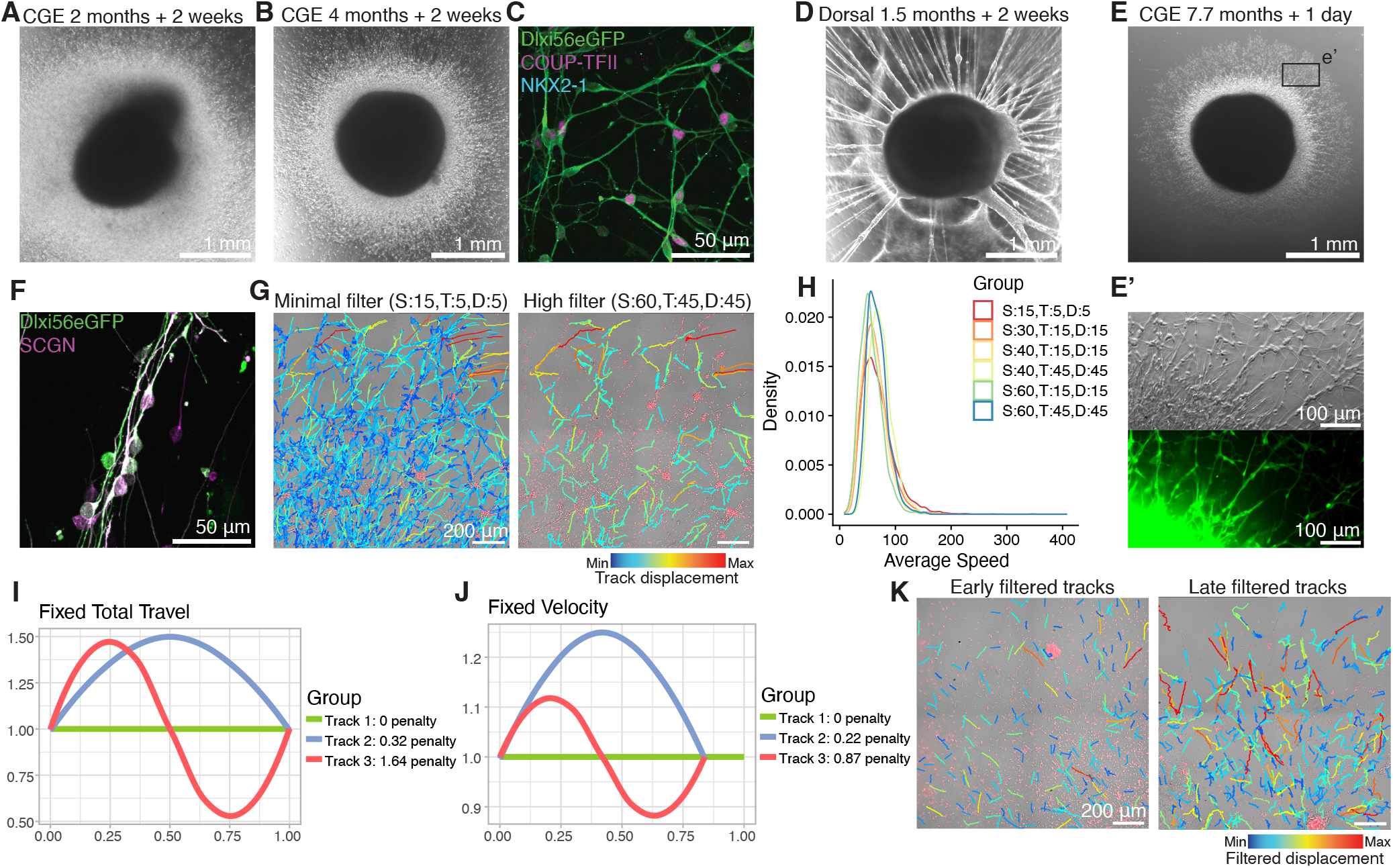
Tracking of early and late CGE INs. **(A)** Brightfield image of a 2-month CGE explant after 2 weeks of plating. **(B)** Brightfield image of a 4-month CGE explant after 2 weeks of plating. **(C)** Immunostaining of 2-month CGE explant show GFP INs are COUP-TFII+. **(D)** Brightfield image of D42/ 1.5-month dorsal patterned organoid explant after 2 weeks of plating. **(E)** Brightfield image of D234/ 7.7-month CGE explant after 1 day of plating. e’, Zoom in of (e) inlet shows INs already migrate from the explant in chains after 1 day. **(F)** IN chains from late CGE explants are SCGN+. **(G)** All identified tracks from one late time-lapse after (left) minimal filter and after (right) stringent filter. S: minimum spots per track, T: minimum travelled distance in pixels, D: minimum net displacement in pixels. Tracks are colored by displacement, showing stringent filter removes very short tracks (dark blue). **(H)** Density plot showing comparisons of the average speed distribution across all tracks with minimal (top/ red) to stringent (bottom/ blue) filtering applied. S: minimum spots per track, T: minimum travelled distance in pixels, D: minimum net displacement in pixels. Stringent filtering (blue) is representative of all tracks. **(I)** and **(J)**, Track examples for the curvature metric (L2 penalty). **(I)** Tracks with an invariant total travel but different velocity. **(J)** Tracks with an invariant fixed velocity but different total travel. In both cases, an increase in curvature leads to a higher penalty. See Methods. **(K)** All identified tracks from an early and late video when same filters (S:60, T:45, D:45) are applied. Late videos have tracks with higher displacement due to faster migration in the same time frame. Tracks are colored by displacement.

**Fig. S5.**
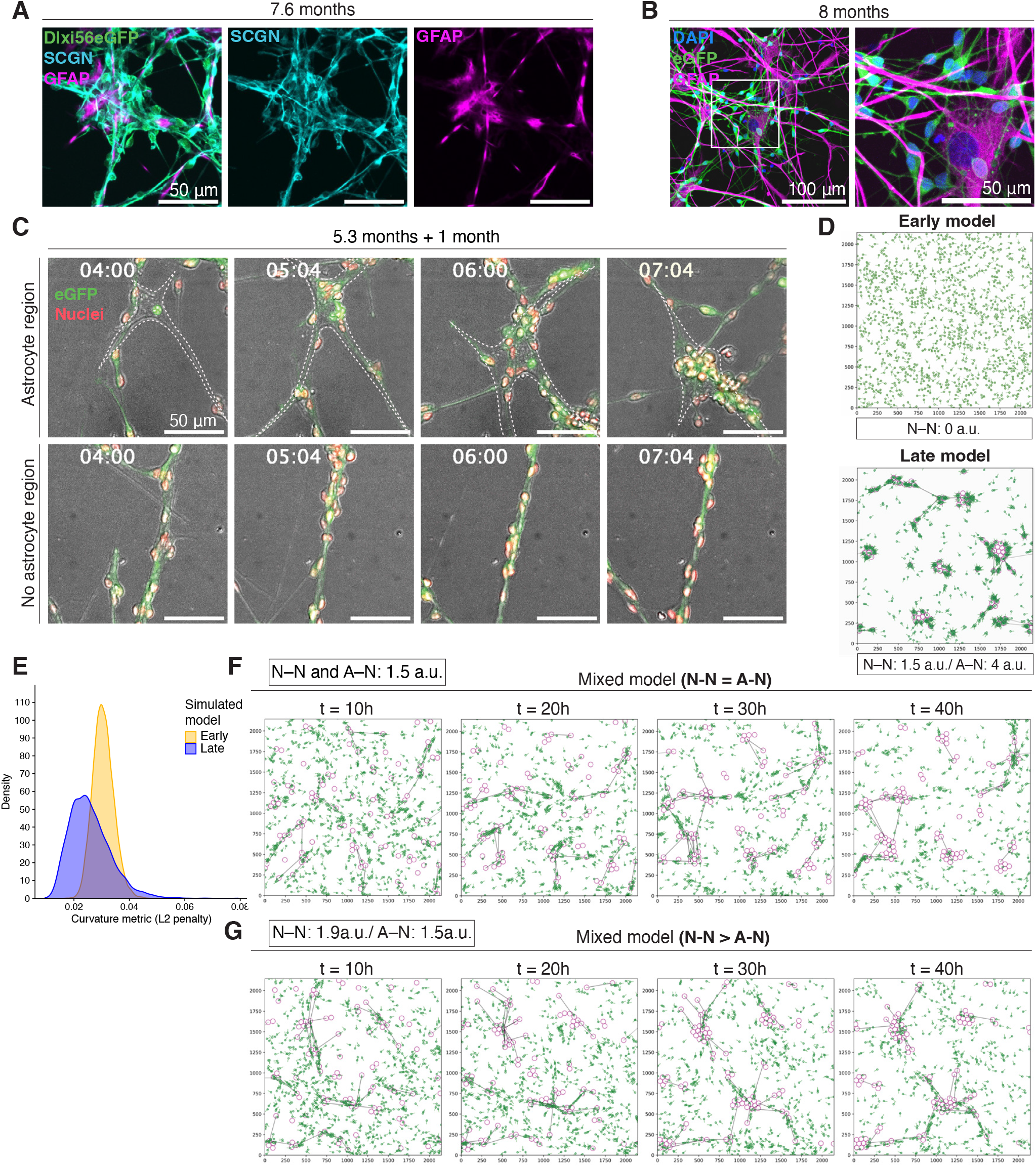
Astrocytes dynamically interact with late-migrating INs. **(A)** 7.6-month explant clusters consist of SCGN+ INs and GFAP+ astrocytes. **(B)** Immunostainings of 8-month explants after 2 weeks of plating show astrocyte and IN morphology and interactions. **(C)** Time-lapse imaging stills of (top) Dlxi56-eGFP+ INs interacting with an astrocyte (white outline) across 3 hours. (Bottom) Time-lapse imaging stills from a different region where INs exhibit chain migration without astrocyte interaction. Imaged with brightfield, GFP, RFP overlay. RFP marks live nuclear tracker dye. D160/ 5.3-month organoid, plated for one month. **(D)** Images at 40 hours of the mathematical simulation of the (top) ‘Early’ condition with no astrocytes and no N-N attraction, and (bottom) ‘Late’ condition with astrocytes present and both N-N and A-N attraction. Astrocytes in magenta, IN in green. **(E)** Density plot showing distribution of track curvature (L2 penalty), comparing simulated early and late conditions. A lower L2 penalty represents straighter tracks. N=6 simulations per condition. **(F)** and **(G)** Images of the mixed model (simMix) simulation across time from 10-40h with altered attraction strengths. **(F)** When N-N and A-N attraction strength is the same, and **(G)** when N-N is stronger than A-N. In both cases, clustering does not occur.

**Fig. S6.**
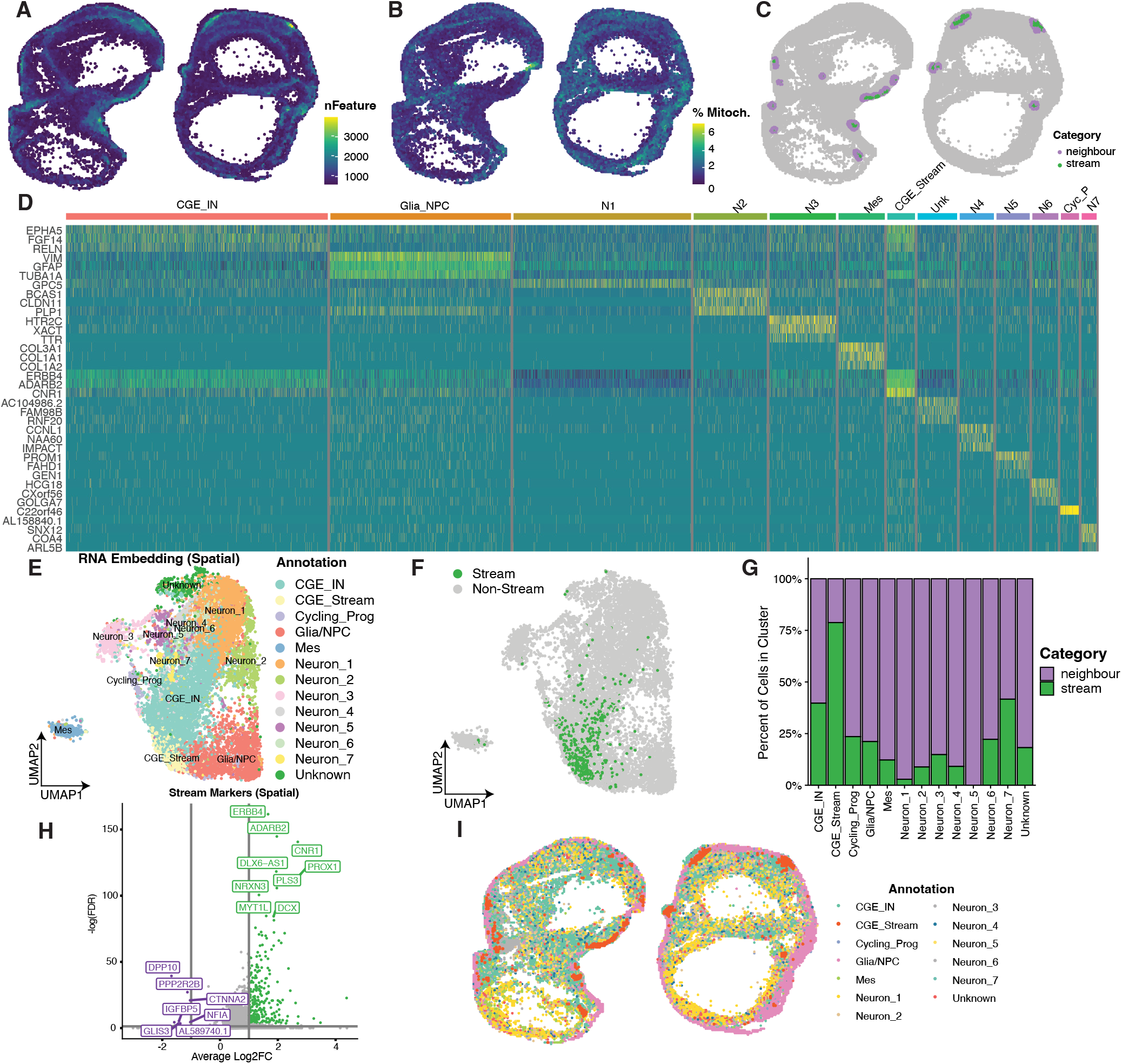
Spatial scRNA-seq in 9-month-old CGE-enriched organoids. **(A-C)**, Spatial embedding of filtered cells from 9 m.o. CGE organoids indicating **(A)** number of features detected per cell, **(B)** percentage of mitochondrial reads per cell, and **(C)** stream membership metadata category. **(D)** Heat map of the top differentially expressed genes per Louvain cluster from spatial scRNA-seq, calculated using Wilcoxon Rank Sum. **(E)** Louvain clustering and UMAP embedding of the gene expression data from spatial scRNA-seq. **(F)** Location of Stream INs in the UMAP embedding. **(G)** Proportional representation of stream INs in each of the clusters. **(H)** Differentially expressed genes marking stream INs compared to other neuronal clusters, calculated with Wilcoxon Rank Sum. **(I)** Spatial embedding of clusters annotated from the transcriptional analyses.

**Fig. S7.**
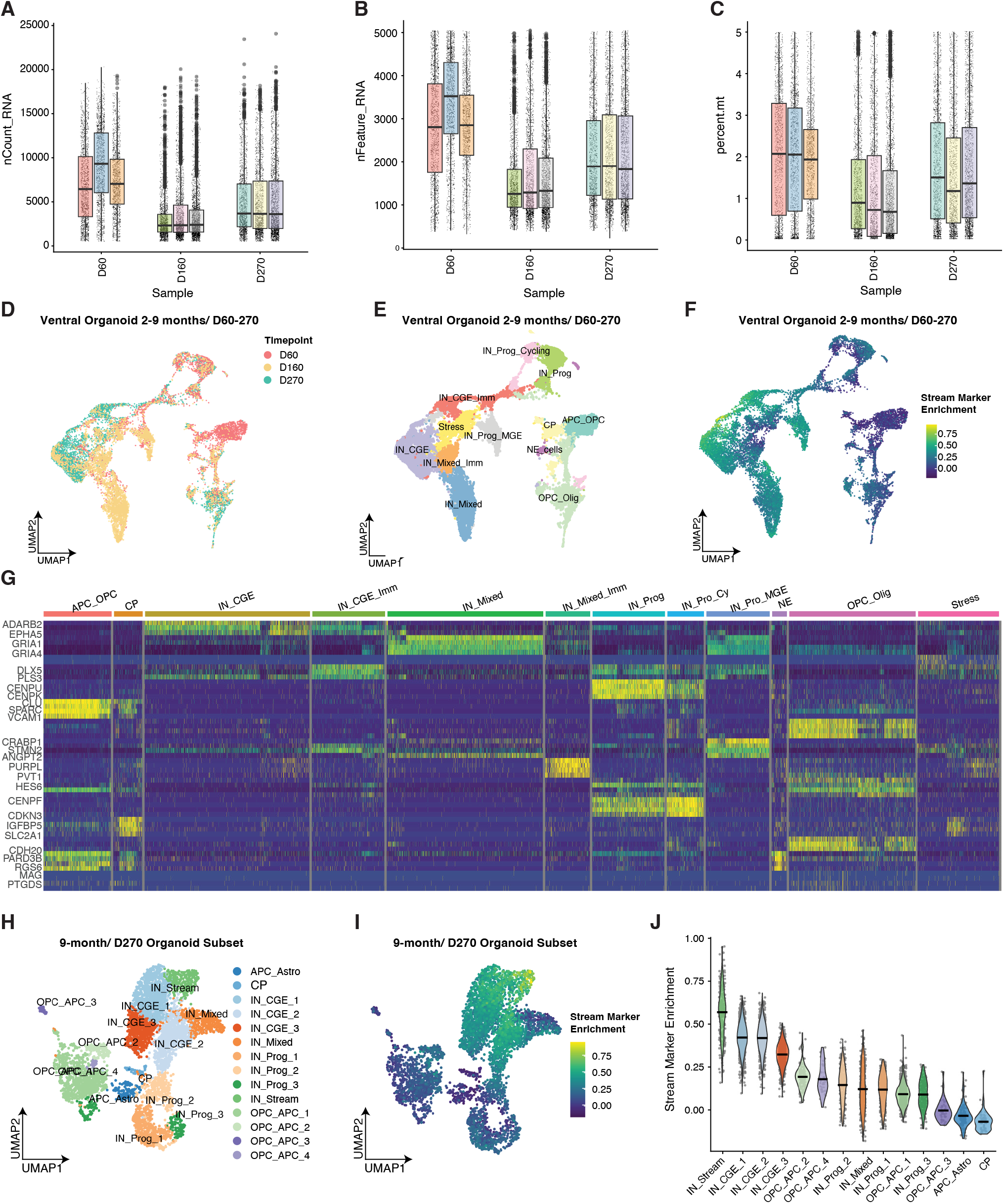
Integrated scRNA-seq in 2-, 5-, and 9-month-old CGE-enriched organoids. **(A-C)** Boxplot of **(A)** raw counts, **(B)** number of detected features, and **(C)** percentage of mitochondrial reads, per timepoint and demultiplexed sample. **(D)** UMAP embedding of harmonized samples across samples, colored by sample timepoint. **(E)** Louvain clusters and annotations of integrated organoid samples. **(F)** Module enrichment score in the integrated samples for the top 200 stream markers genes identified from the spatial scRNA-seq. **(G)** Heatmap of the top marker genes per Louvain cluster in the integrated scRNA-seq data, calculated by Wilcoxon Rank Sum. **(H)** UMAP embedding and higher-resolution Louvain clustering of the subset of samples from the 9-month timepoint. **(I)** Module enrichment score in the 9 m.o. samples for the top 200 stream markers genes identified from the spatial scRNA-seq. **(J)** Violin plot of stream marker enrichment per higher-resolution cluster in 9 m.o. CGE-enriched organoids.

**Fig. S8.**
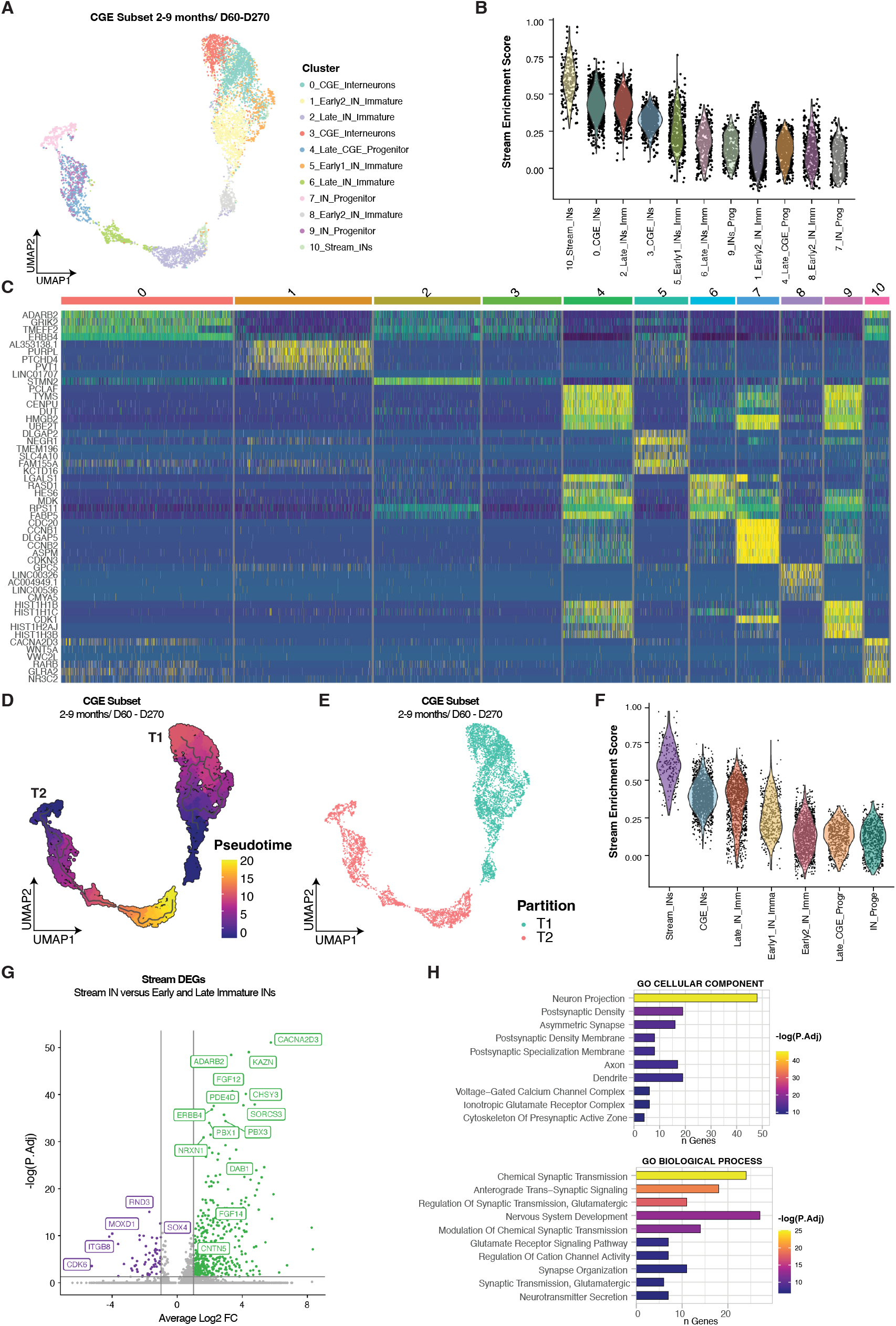
Integrated CGE trajectory analysis. **(A)** The CGE trajectories identified by Monocle3. The CGE lineages were isolated from the integrated scRNA-seq object through successive subsets based on canonical CGE marker expression. These cells were aligned by sample, and clusters were identified and annotated using the previously defined cluster labels. **(B)** Stream marker enrichment scores per cluster in the CGE trajectories. **(C)** Heatmap of the top marker genes per Louvain cluster in the CGE trajectories, calculated by Wilcoxon Rank Sum. **(D)** Pseudo-time scores of the CGE trajectories identified by Monocle 3. **(E)** Trajectory partitions identified by Monocle3. **(F)** Stream marker gene enrichment scores by cell type annotation in the CGE trajectories. **(G)** Differentially expressed genes in the Stream IN cluster compared to the other CGE IN clusters. **(H)** Gene Ontology enrichments for DEGs positively marking the Stream INs, calculated using EnrichR for GO Cellular Component and GO Biological Process.

**Fig. S9.**
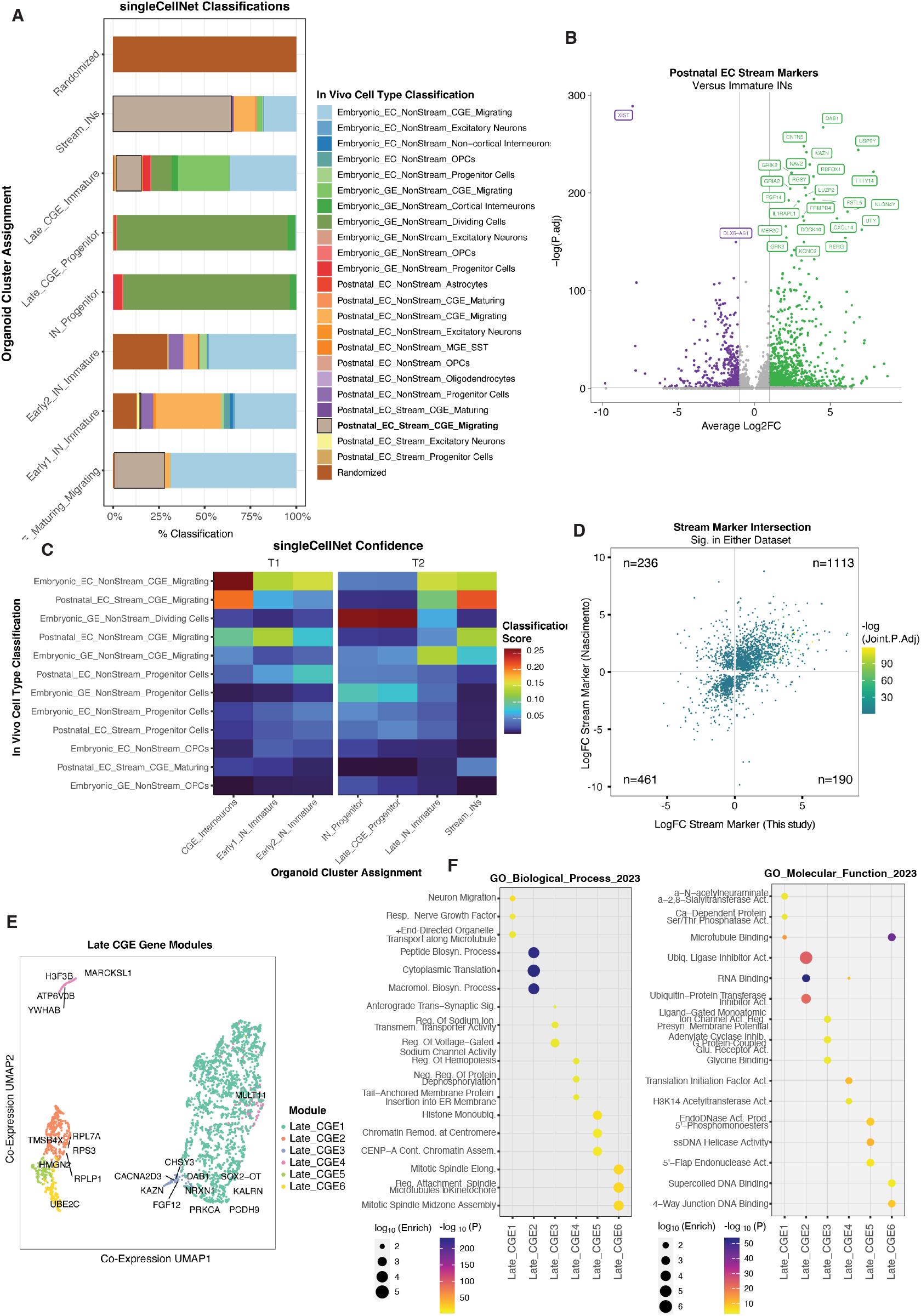
Comparison of scRNA-seq to *in vivo* datasets and gene module analysis. **(A)** Full singleCellNet assignment proportions for each of the CGE organoid clusters to cell populations present in Nascimento et al, 2024. The classifiers were trained on the labels present in the legend; labels were generated for *in vivo* cell types by intersecting timepoint and cell type metadata and comparing marker gene expression. **(B)** Differentially expressed genes in the Postnatal EC Stream cluster compared to the other immature INs from the *in vivo* dataset. **(C)** Heatmap of mean classification scores identified by singleCellNet for each CGE organoid cluster based on its training for the labels in **(A)**. Organoid clusters are faceted by trajectory. **(D)** Intersection of shared EC stream and organoid stream markers significant in at least one dataset. EC Stream Markers were recalculated using Wilcoxon Rank Sum of EC Stream CGE INs compared to other immature CGE INs in the *in vivo* dataset to generate a set of markers. **(E)** UMAP embedding of genes in modules based on expression across cells the integrated late CGE lineage. Each gene (point) is colored by module membership identified by hdWGCNA. **(F)** Gene ontology enrichments were calculated for each module identified by hdWGCNA by Biological Process and Molecular Function.

**Fig. S10.**
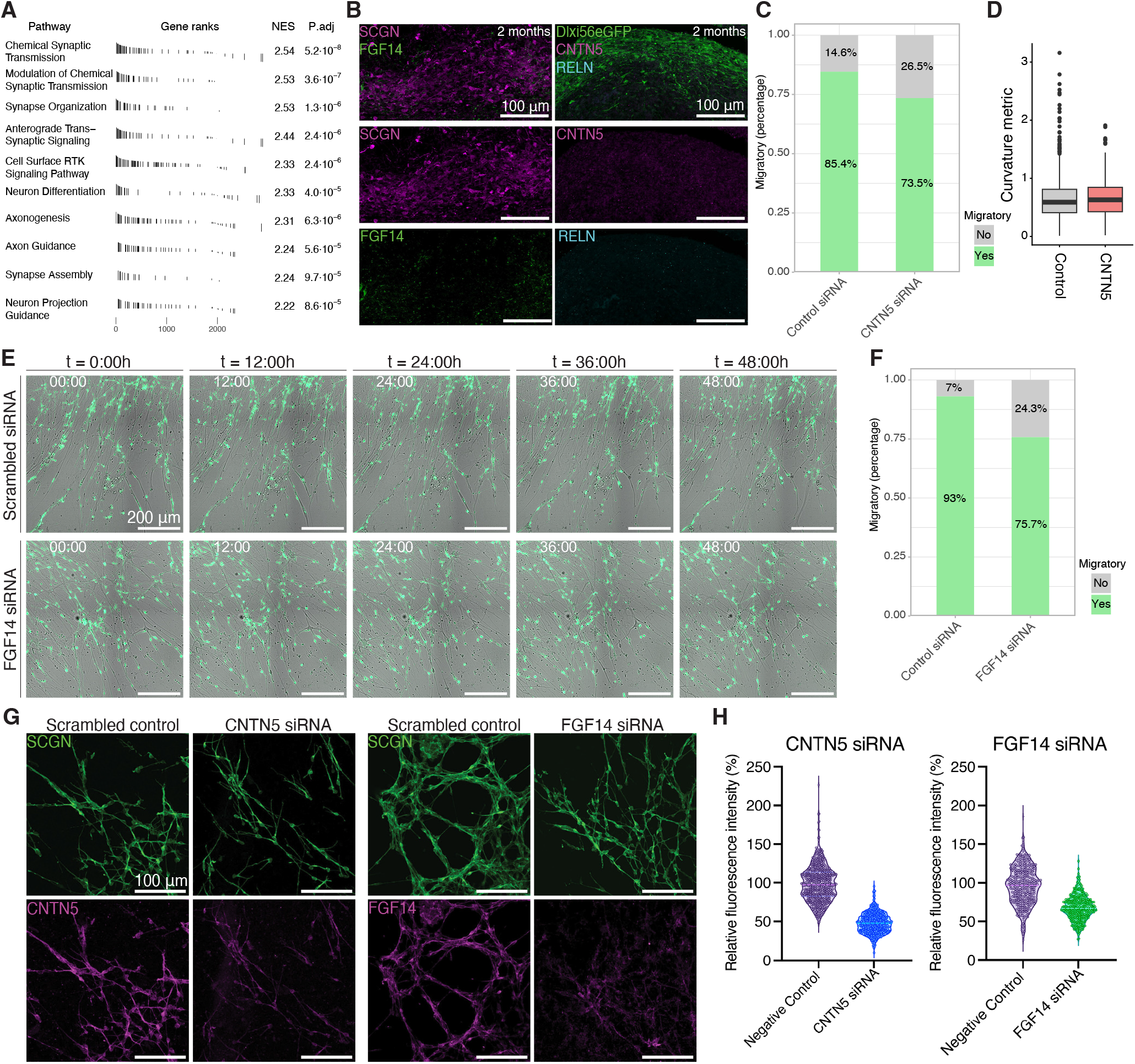
Transient knockdown of *FGF14* and *CNTN5* in late explants. **(A)** GSEA analysis on module 3 genes for GO terms. **(B)** Immunostainings of 2-month organoids for (left) *FGF14* and SCGN, and (right) *CNTN5* and RELN. **(C)** Stacked bar plot showing percentage of migratory (green) vs. non-migratory (grey) INs after control siRNA and *CNTN5* siRNA. N=6 for control, N=8 for *CNTN5*. **(D)** Box plot showing curvature metric/ L2 penalty of highly migratory tracks comparing control and *CNTN5* KD. *CNTN5* tracks are not significantly different from control after filtering at p = 0.372648234. **(E)** Stills every 12h from a 48h time-lapse imaging video after 10-month explant KD with (top) scrambled control and (bottom) *FGF14* siRNA. **(F)** Stacked bar plot showing percentage of migratory (green) vs. non-migratory (grey) INs after control siRNA and *FGF14* siRNA. N=6 for control, N=8 for *FGF14*. **(G)** Immunostainings after 2 weeks of plating for (left) SCGN and *CNTN5*, and (right) SCGN and *FGF14* in control and siRNA conditions. siRNA conditions show reduced fluorescence intensity of both *CNTN5* and *FGF14*. **(H)** Quantifications of fluorescence intensity for *CNTN5* and *FGF14* immunostainings in negative control and with KD. Fluorescence intensity is set relative to control as a percentage. N=6 for *CNTN5*, N=5 for *FGF14*.

## Supplementary Movie Legends

**Movie S1**

Chain migration in organoid streams. Video of Fig. 2I, J. Chain migration of a stream in 8.7-month whole CGE-Os. GFP marks Dlx5/6-interneurons. 7 fps. Time stamp in hours. Scale bar = 100 *µ*m.

**Movie S2**

Early organoid explant migration. Video of Fig. 3C (top). Dlxi56-eGFP+ interneurons migrate out of young (Day 60 plated) explants as individual, dispersed neurons. Images acquired every 20 mins. Time stamp in hours. Scale bar = 100 *µ*m.

**Movie S3**

Late organoid explant migration. Video of Fig. 3C (bottom). Dlxi56-eGFP+ interneurons migrate out of old (D160 plated) explants in connected chains and form large clusters. Images acquired every 20 mins. Time stamp in hours. Scale bar = 100 *µ*m.

**Movie S4**

Early organoid explant tracking. Video of Fig. 3f (top). Interneurons migrating out of a young (3 months/ D90) explant, tracked with red nuclear dye and visualized in brightfield. Images acquired every 2 minutes for tracking. Time stamp in hours. Scale bar = 100 *µ*m.

**Movie S5**

Late organoid explant tracking. Video of Fig. 3f (bottom). Interneurons migrating out of an old (7 months/ D223) explant, tracked with red nuclear dye and visualized in brightfield. Images acquired every 2 minutes for tracking. Time stamp in hours. Scale bar = 100 *µ*m.

**Movie S6**

Interneurons interact with astrocytes in late explants. Video of fig. S5C (top). After 1 month of plating, GFP+ interneurons in 6.3-month explants interact with unlabeled astrocytes in some regions. Scale bar = 100 *µ*m.

**Movie S7**

Interneurons migrate in chains without astrocytes in late explants. Video of fig. S5C (bottom). After 1 month of plating, GFP+ interneurons in 6.3-month explants maintain independent chain migration in regions with low astrocyte density. Scale bar = 100 *µ*m.

**Movie S8**

Homophilic model (simHomo) N-N=1.5. Simulation of the homophilic model shown in Fig. 4J. Astrocytes marked by magenta circles, interneurons marked by green dots. Interneurons spontaneously form chains. T=20 min intervals. Total simulation time = 40h

**Movie S9**

Hetereophilic model (simHetero) A-N=4. Simulation of the heterophilic model shown in Fig. 4K. Astrocytes marked by magenta circles, interneurons marked by green dots. Chains and clusters do not form. T=20 min intervals. Total simulation time = 40h

**Movie S10**

Mixed model (simMix) N-N=1.5, A-N=4. Simulation of the mixed model shown in Fig. 4L. Astrocytes marked by magenta circles, interneurons marked by green dots. Both N-N and A-N attraction leads to formation of streams and clusters, as seen in explant assays. T=20 min intervals. Total simulation time = 40h

